# Discovery of hypercompact epigenetic modulators for persistent CRISPR-mediated gene activation

**DOI:** 10.1101/2023.06.02.543492

**Authors:** Giovanni A. Carosso, Robin W. Yeo, T. Blair Gainous, M. Zaki Jawaid, Xiao Yang, James Y.S. Kim, Kavita Jadhav, Nina Juan-Sing, Siddaraju V. Boregowda, Vincent Cutillas, Lei Stanley Qi, Alexandra Collin de l’Hortet, Timothy P. Daley, Daniel O. Hart

**Affiliations:** Epicrispr Biotechnologies, South San Francisco, CA 94080, USA; Department of Bioengineering, Stanford University, Stanford, CA 94305, USA; Sarafan ChEM-H, Stanford University, Stanford, CA 94305, USA; Chan Zuckerberg Biohub-San Francisco, San Francisco, CA 94158, USA

## Abstract

Programmable epigenetic modulators provide a powerful toolkit for controlling gene expression in novel therapeutic applications, but recent discovery efforts have primarily selected for potency of effect rather than contextual robustness or durability thereof. Current CRISPR-based tools are further limited by large cargo sizes that impede clinical delivery and, in gene activation contexts, by brief activity windows that preclude transient, single-dose strategies such as lipid nanoparticle (LNP) delivery. To address these limitations, we perform high-throughput screening to discover novel classes of transcriptional modulators derived from thousands of human, viral, and archaeal proteomes. We identify high-potency activators capable of mitotically stable gene activation in a multitude of cellular contexts and leverage machine learning models to rationally engineer variants with improved activities. In liver and T-cells, novel hypercompact activators (64 to 98 amino acids) derived from vIRF2 core domain (vCD) achieve superior potency and durable activation lasting weeks beyond the current large activators (∼five-fold larger). In a humanized mouse model, we target a human hypercholesterolemia susceptibility gene and achieve activation persisting five weeks after a single dose by LNP delivery. Our discovery pipeline provides a predictive rubric for the development of contextually robust, potent, and persistent activators of compact size, broadly advancing the therapeutic potential of epigenetic gene activation.

## Introduction

RNA-guided epigenetic control of gene expression via nuclease-dead CRISPR/Cas (dCas) systems offers potential therapeutic avenues for treating a wide range of disease indications without creating double strand breaks or changing the DNA sequence^1-3^. However, existing tools are limited in clinical utility by large coding DNA cargo sizes, variable efficacies at diverse targets, and temporally transient windows of activity^4,5^. Traditional approaches to programmable target gene activation have relied on a limited catalog of annotated peptide domains sourced from eukaryotic and viral proteomes, or fusions thereof^6-8^. Such domains act by recruiting transcriptional cofactors in part through acidic and hydrophobic contact interfaces^9-11^, or by encoding enzymatic domains that alter DNA methylation status or histone post-translational modifications^12-14^. Emerging literature suggests highly disparate activity levels among these modulatory mechanisms in terms of potency, context-dependent robustness, and durability of effect^15,16^. Expanding the set of tools available for transcriptional modulation is therefore critical to achieve control of target gene expression in distinct chromatin contexts, particularly in therapeutic applications where a single-dose, transient vector such as lipid nanoparticle (LNP) delivery may be favored over AAV vectors. Unbiased tiling and testing of diverse proteomes with high-throughput screening is a powerful method for identifying compact peptide fragments capable of transcriptional modulation^17-19^. The most potent peptide fragments, in turn, can provide a basis for iterative refinement of desirable modulator properties by subsequent rounds of rational engineering^20-22^.

While the human genome encodes over 2,000 transcription factors and chromatin regulators^23^, virological surveys estimate there exists up to 320,000 mammalian-tropic viral species among the estimated 10^31^ viral particles comprising the global virome^24,25^. Key to viral evolution is their ability to manipulate host gene expression, suggesting that these genomes represent a vast reservoir of rapidly evolving molecular tools optimized for transcriptional modulation with compact coding size^26^. Previous studies have separately evaluated sets of either human- or viral-encoded domains^27,28^, but direct comparisons of their transcriptional functions at multiple endogenous human targets remain to be performed. Additionally, the restrictive environmental constraints of other biological classes, such as extremophilic archaea, represent a previously underexplored resource for transcriptional tiling, and could contain peptide domains with unique biochemical properties and functionalities^29^.

Unmodified histone tails carry a positive charge that is believed to drive electrostatic interactions with DNA and neighboring nucleosomes contributing to a compacted chromatin environment. This in turn restricts engagement by the transcriptional apparatus, preventing gene expression in the default state^30^. While repressive epigenetic marks such as DNA methylation and H3K27me3/H3K9me3 marks on histone tails can be propagated heritably during cellular replication, epigenetic persistence of permissive chromatin marks remains to be determined ^31-36^. Accordingly, while others have demonstrated mitotically durable gene silencing via dCas-mediated transcriptional modulation, gene activation has thus far shown a limited activity window with transcriptional upregulation dissipating in the absence of the initiator^8,37-39^. Prolonged, though still transient, activation windows have been reported via co-deposition of active marks H3K4me3 and H3K79me3 by dCas9 fusions to PRDM9 and DOT1L histone methyltransferases, respectively, or via DNA demethylation by dCas9-Tet1, suggesting that opportunities exist for mitotically stable gene activation^13,14^. Intrinsic programs exist for cells to durably activate gene expression programs, such as in early embryonic development, that persist heritably despite the challenges of rapid cell division and differentiation^40-42^. Moreover, cellular environmental responses require sustained gene activation, as in H3K4me1-mediated p65 induction following transient hyperglycemic spike in aortic endothelial cells^43^, or during innate immune responses with H3K27ac-mediated immune gene activation which precedes DNA demethylation^44,45^. Mitotically stable activation memory, if mediated by H3K27ac, would likely require cognate epigenetic readers, such as CBP/P300-associated BET family bromodomains, to recognize and redeposit the permissive acetyl marks on nucleosomes via ‘read-write’ mechanisms^46,47^.

Here, we performed high-throughput dCas-mediated recruitment screens to systematically interrogate the transcriptional potencies of tens of thousands of human, viral, and archaeal-derived peptide sequences in both silenced and expressed promoter contexts; we identified known and novel peptide domains with strong biases toward activation by viral domains, suppression by human domains, and context-dependent dual-activity profiles for archaeal domains. Sequence-based predictions, paired with extensive validation testing in diverse chromatin contexts, revealed a predictive biochemical rubric for engineering functional improvements to minimal core activator domains. Importantly, a subset of these activators displays a novel ability to maintain target activation through dozens of cell divisions after a single transient delivery in multiple contexts, and ultimately maintained hepatic gene activation of a therapeutically relevant human gene, low-density lipoprotein receptor (*LDLR*), *in vivo* at five weeks after a single-dose LNP administration. This pipeline for modulator discovery and engineering yielded hypercompact transcriptional activators as small as 64 amino acids (a.a.) that outperform previous benchmarks with respect to potency, context-independent robustness, and mitotic durability of effect, despite occupying ∼12-20% of their protein-coding cargo size.

## Results

### Identification of transcriptional modulator domains by high-throughput screening

To interrogate the transcriptional potency of naturally occurring peptide fragments, we first generated libraries by tiling across candidate human, viral, and archaeal genomic coding sequences (**Fig. 1a** and **Extended Data Fig. 1a**). The human set consisted of human nuclear-localized proteins (n=549) including DNA- and histone-modifying enzymes, chromatin remodelers, and transcription factors. We then curated a custom set of viral genome coding sequences (n=3,548) distributed across 188 viral families. Selections were weighted to enrich for viruses that evolved predominantly in mammalian host reservoirs or have undergone zoonotic transfer, reasoning that such domains may be adapted to modulate human gene expression^48,49^. We further sought to ask whether domains from acidophilic, thermophilic, or otherwise extremophilic viruses and host archaea might confer more robust or heritable transcription-modulatory functions, owing to evolutionary pressures that favor vertical versus horizontal host transmission, so we included viral peptides from metagenomics surveys (n=129)^50,51^ and the full proteome of volcanic archaeon *Acidianus infernus*^52^. Lastly, we added a set of annotated human virus transcriptional regulators (vTRs)^26^ (n=419). As positive controls, we added human proteins with known transcriptional function, and GC-matched random-sequence negative controls. We partitioned the sequences into overlapping DNA tiles encoding 85-a.a. peptide fragments and pooled these into a merged screening library of uniquely barcoded oligonucleotides (n=43,938) to enable group-wise comparisons of modulator potency.

**Fig. 1.**
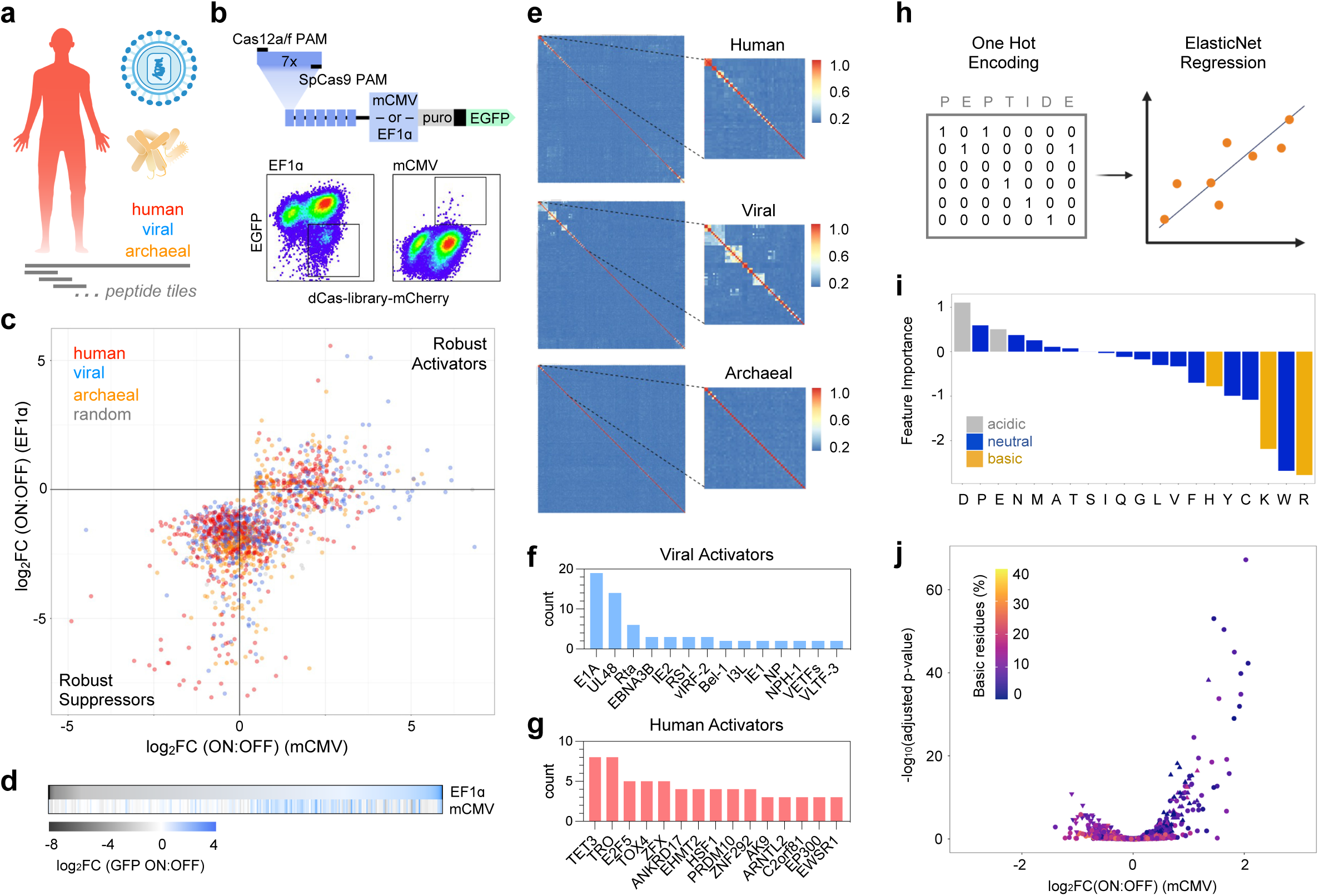
Identification of transcriptional modulator domains by high-throughput screening. **a,** Schematic depiction of pooled human, viral, and archaeal coding libraries and peptide tiling strategy. **b,** Schematic depiction of custom mCMV-GFP and EF1ɑ-GFP promoters targeted for pooled modulator screening. Representative FACS plots of EF1ɑ-GFP (left) and mCMV-GFP (right) K562 cells following lentiviral transduction of the pooled dCas9-library fusions. boxes represent OFF and ON gates for EF1ɑ-GFP and mCMV-GFP cell lines respectively. **c,** Scatterplot comparing activation strengths (log_2_ fold-change enrichment of barcode counts in ON vs OFF bins) of top-significance modulators (n=560 activators, n=1,330 suppressors) from mCMV-GFP and EF1ɑ-GFP screens respectively. Robust activators, suppressors, and dual activity hit quadrants are indicated. **d,** Heatmap of activation strengths (log_2_ fold-change enrichment of barcode counts in ON vs OFF bins) for statistically significant modulators indicating activation versus suppression activities at mCMV-GFP and EF1ɑ-GFP promoters. **e,** Sequence homology clustering of human (top, n=209), viral (middle, n=177), and archaeal (bottom, n=154) activators identified in the mCMV-GFP high-throughput screen identifies clusters of modulators with sequence similarity. **f,g,** Ranked lists of viral (**f**) and human (**g**) genes from which statistically significant activation tiles originate. **h,** Schematic depicting featurization of peptide sequences by OneHot encoding and subsequent training of a generalized linear regression model (ElasticNet) to predict activation strength based on peptide sequence alone. **i,** Extracted feature importance from top generalized linear regression model detailing which residue types were predictive of activation strength (gray: acidic, blue: neutral, gold: basic). **j,** Volcano plot illustrating screening results from the mCMV-GFP validation screen colored by composition of basic residues. Validation tiles are represented by circles, positive controls (activation) by triangles, and positive controls (suppression) by inverted triangles.

We engineered a pair of GFP reporter systems driven by either a minimal CMV (mCMV) promoter (OFF-to-ON) or EF1ɑ promoter (ON-to-OFF) to screen for activators and suppressors, respectively (**Fig. 1b**,top and **Extended Data Fig. 1b**). As control, the mCMV-GFP cell line could be activated by dCas9-VPR and the EF1ɑ-GFP cell line could be suppressed by dCas9-KRAB (**Extended Data Fig. 1c**). mCMV-GFP and EF1ɑ-GFP reporter cells stably expressing a targeting Cas9 sgRNA were lentivirally transduced with dCas9-modulator library fusions, then sampled by FACS-based separation of GFP-ON and GFP-OFF cells (**Fig. 1b**,bottom). We computed barcode enrichments in GFP-ON versus GFP-OFF cells, identifying 560 activators and 1,330 suppressors of mCMV-GFP and EF1ɑ-GFP, respectively (**Fig. 1c** and **Extended Data Fig. 1d-f**) with taxon-dependent potencies (**Extended Data Fig. 1g**). Comparisons between both screens provided early insights into contextual robustness; among the 560 mCMV-GFP activators, 303 were depleted among EF1ɑ-GFP suppressors (“robust activators”), i.e. concordance of effect at both promoters (**Fig. 1c,d** and **Extended Data Fig. 1h**). Similarly, among 1,330 EF1ɑ-GFP suppressors, 622 were depleted among mCMV-GFP activators (“robust suppressors”). Random negative control tiles showed a uniform distribution about log_2_FC=0 in both screens as expected (**Extended Data Fig. 1i**).

While activator hits were distributed proportionately across viral (n=177), human (n=209), and archaeal (n=154) tiles, the most potent were viral in origin and segregated into distinct sequence homology clusters (**Fig. 1e,f, Extended Data Fig. 1j**) of both known and novel classes: E1A (hAdv), VP16 (HHV), VLTF3 (Pox), IE1/IE2 (HCMV), and vIRF2 (HHV). Human activator clusters largely consisted of homologous tiles from different isoforms of major families including E2F5, ARNTL2, ZFX/ZFY, NHSL1, TET3, HSF1, MESP1, and TOX4 (**Fig. 1e,g**, **Extended Data Fig. 1j**). We next trained a regularized regression model to predict activation strength based on peptide sequence (**Fig. 1h**) and extracted scores for amino acid importance (**Fig. 1i** and **Extended Data Fig. 1k,l**). While enrichment of acidic residues is indeed predictive of activation, as previously reported^11^, we find that depletion of basic residues is a far stronger biochemical predictor (**Fig. 1j, Extended Data Fig. 1m**). Interestingly, activators display taxonomy-dependent biochemical trends; viral activators are strongly enriched for acidic residues while human activators are primarily depleted of basic residues (**Extended Data Fig. 1k**). To further investigate biochemical and biophysical design rules, we generated 3-dimensional structure predictions using ESMFold and computed quantitative structural features using Yasara (see Methods)^53,54^, and found that activator sequences are enriched for small amino acids, intolerant of ꞵ-sheets, and enriched for turns (**Extended Data Fig. 1m**). In contrast to previous reports^10,11^, we do not observe enrichment of hydrophobic residues, suggesting that they are dispensable for activation of this locus (**Extended Data Fig. 1m,n**). Results from a subsequent hit sub-library screen (**Extended Data Fig. 2a-j**) independently validated our top hits and were consistent with these biochemical predictions.

Suppressor hits were dominated by human tiles in potency, followed by archaeal tiles, although overall hit frequencies were once again distributed proportionately across human (n=502), viral (n=388), and archaeal (n=389) origins (**Fig. 1c** and **Extended Data Fig. 1e**). Human ZNF-derived tiles were the most potent suppressors, clustering based on KRAB-like domains or ZF C2H2 domains (**Extended Data Fig. 3a-c)**. Surprisingly, hydrophobic residue enrichment was greater among suppressors than activators (**Extended Data Fig. 3d)**. Archaeal suppressors, while poorly annotated, displayed higher potencies with increasing content of aliphatic residues and ꞵ-sheets (**Extended Data Fig. 3e**). Both activators and suppressors exhibited a preference for a smaller solvent-accessible surface area (SASA) and presence of charged, acidic residues conferring negative electrostatic potential (**Extended Data Fig. 3f**).

### Biochemical features predict contextually robust activator domains

We next focused on *in-silico* predictions of robustness, i.e. context-independent potency, among the less characterized or unannotated viral and archaeal elements, based on amino acid composition. Relative to null controls, robust viral activators are depleted of basic and aromatic residues and ꞵ-sheets, and contain fewer aliphatic, polar, and hydrophobic residues and lower helix and flexibility (B-factor) scores (**Extended Data Fig. 1k-n**). Further, they have highly negative net charge and electrostatic potential, low mass, prefer higher content of small, acidic, and coil residues, and exhibit a non-zero degree of turn residues. Robust viral suppressors display fewer biochemical biases beyond hydrophobicity but are weakly enriched for charged and acidic residues and more tolerant to ꞵ-sheets, while robust archaeal suppressor potency increases with aliphatic residues and ꞵ-sheets (**Extended Data Fig. 3c-f**).

To test these predictions in a context-dependent manner, we designed sgRNAs targeting multiple human and synthetic promoters and evaluated modulator potencies by recruitment via dCasMINI^55^ as well as dCas9. We selected a set of viral and archaeal modulators (n=95) with variable potency predictions in our K562 cell screens (**Extended Data Fig. 1e,f**) and diverse biochemical features (**Extended Data Fig. 1n**), and compared their potencies across promoter contexts in HEK293T cells relative to canonical benchmarks p65, Rta, VP64, tripartite VP64-p65-Rta (VPR), and ZNF10 KRAB by measuring induced expression changes at protein and mRNA level (**Fig. 2a** and **Extended Data Fig. 4, Extended Data Fig. 5**). We observe viral activators that match or outperform many benchmarks, except for VPR in most cases, in potency (**Fig. 2a-c**, **Extended Data Fig. 4**, **Extended Data Fig. 5**). Overall concordance of target effects was greatest for the predicted robust viral activators, independently of dCas recruitment system (**Extended Fig. 4a,b**). Viral and archaeal suppressors outperformed human ZNF10 KRAB in some cases (**Fig. 2a-c**, **Extended Data Fig. 4**, **Extended Data Fig. 5**), but overall lacked the contextual robustness of the benchmark. Consistently, the most robust viral activators were those most depleted of basic residues (**Fig. 2d,e**), lacked ꞵ-sheets, and were less enriched for hydrophobic residues than suppressors (**Fig. 2f,g**). Robust archaeal suppressors displayed an aliphatic dependence not seen for viral suppressors (**Fig. 2g**). Global comparison of target effects highlighted two high-potency peptides originating from human herpesvirus 8 (KSHV) protein (Q2HR71), both fulfilling our activator rubric of basic residue depletion, low negative charge, low hydrophobicity scores, and other features (**Fig. 2h** and **Extended Data Fig. 6a-e**). We therefore decided to further investigate these tiles and hypothesized that exploiting this set of properties by domain engineering could further enhance transcriptional activation.

**Fig. 2.**
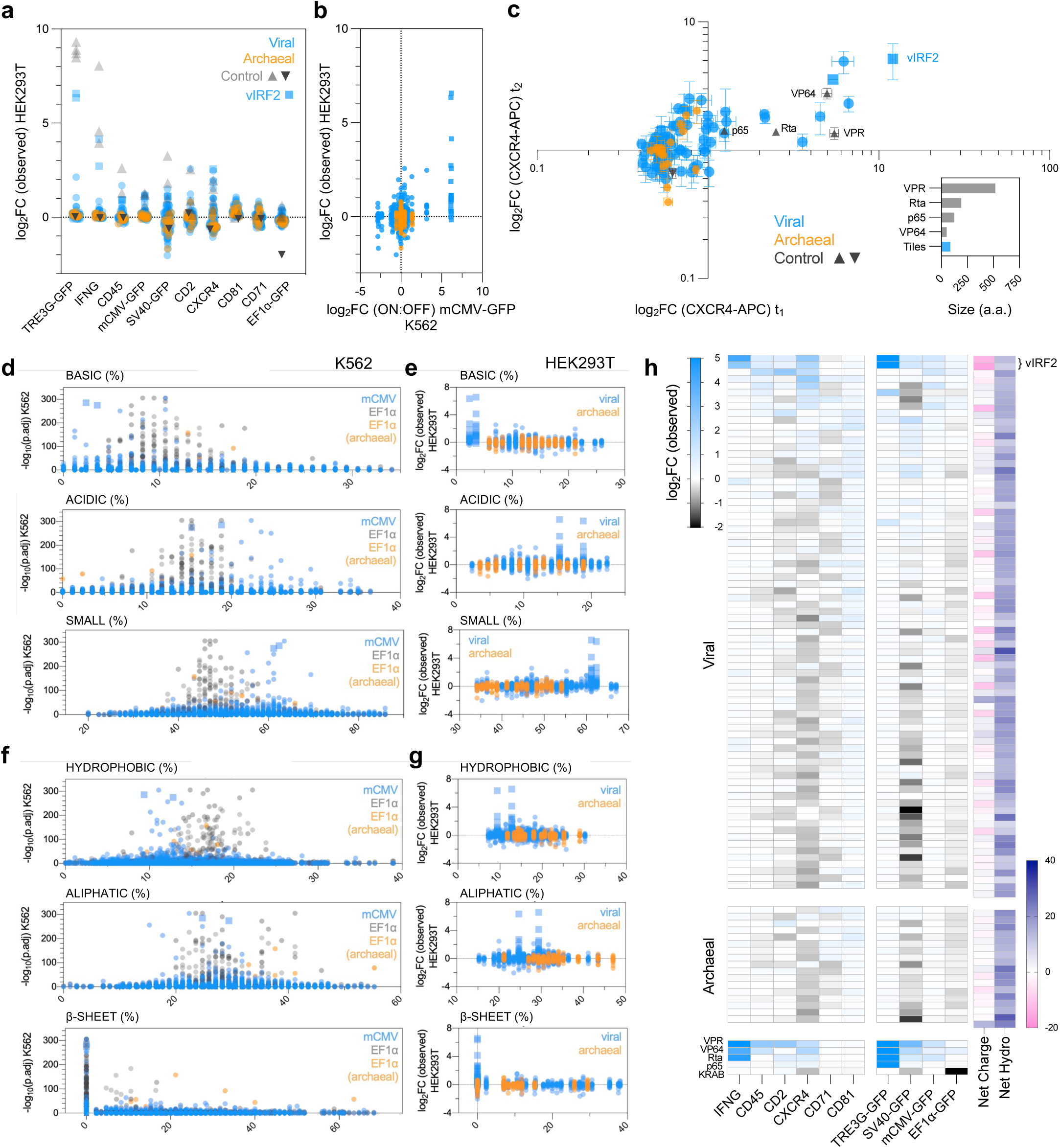
Biochemical features predict robust and context-independent activator domains. **a,** Functional testing by arrayed dCasMINI-mediated recruitment of selected modulator screen tiles (n=95) and benchmark modulators at ten target genes in HEK293T cells. Dots represent observed modulator activity per target (mean fold-change of 3 or more replicates) in observed protein expression, relative to the mean of non-targeting sgRNA (sgNT) and dCasMINI recruited without modulator fusion (dCasMINI) conditions (**Extended Data Fig. 5**). Viral (blue), archaeal (orange), benchmark controls (triangle), and top viral hit tiles (square) are indicated. **b,** Comparison of observed mean activities in HEK293T cells (y-axis) against computed barcode enrichment scores (mCMV-GFP ON:OFF) from pooled high-throughput screens in K562 cells (x-axis) (Fig. 1c,d). **c,** Representative functional testing experiment by flow cytometry at one of the ten targeted contexts shown in (**a**). Fold-changes in CXCR4-APC fluorescence (geometric mean) at 3 days post-transfection (d.p.t.) (t_1_, x-axis) versus 7 d.p.t. (t_2_, y-axis) by each of 95 viral and archaeal modulators and benchmark controls VPR, VP64, Rta, p65, KRAB (mean±SEM). Relative modulator peptide sizes in amino acids (a.a.) plotted for reference (inlaid). **d,e,** Biochemical feature scores (x-axis) plotted against hit significance (ON:OFF, adj. p-val) for the full modulator libraries in mCMV-GFP and EF1ɑ-GFP K562 screens (**d**) or against observed activities in HEK293T cells (mean fold-change per target) (**e**). Features predicting activators are shown. **f,g,** Biochemical feature scores (x-axis) plotted against hit significance (ON:OFF, adj. p-val) for the full modulator libraries in mCMV-GFP and EF1ɑ-GFP K562 screens (**f**) or against observed activities in HEK293T cells (mean fold-change per target) (**g**). Features predicting suppressors are shown. **h,** Ranked heatmap summary of context robustness in modulator activities in arrayed HEK293T cell experiments, with observed protein fold-changes per target (left) and scores for net charge and hydrophobicity (right). Gradients of mean observed activation (blue), suppression (black), and no effect (white) by each modulator at each locus are indicated.

### Engineered potency of vIRF2 core domain (vCD)-based modulators

Positional mapping of our screen tiles revealed an overlap of two exceptionally high-potency and context-robust activators from the viral interferon regulatory factor 2 (vIRF2) KSHV protein (Q2HR71), homologous to human interferon regulatory factors^56^ (**Fig. 3a-c**). Among the 95 characterized viral and archaeal tiles, both tiles were biochemical outliers with respect to basic residue depletion, negative net charge, and low mass (**Fig. 2d,h; Extended Data Fig. 6**). We applied a deep learning model (ADPred)^57^ to identify which residues were predicted to contribute to transactivation domains within the two vIRF2 tiles and identified a 32-a.a. vIRF2 core domain (vCD) located within the overlapping region, suggesting that this vIRF2 core domain may be responsible for the strong activation ability of the vIRF2 tiles (**Fig. 3c**). Despite sharing 68.2% sequence identity, these peptides differed notably in electrostatic potential and turn scores (**Extended Data Fig. 6**), suggesting a role for domain positional effects on secondary structure. We therefore rationally designed a set of vCD-based variants (**Fig. 3d-i, Extended Data Fig. 7a,b**) to test configurations of 1) the isolated core domain by itself in various positions (n=24), 2) double-vCD tandem repeats (n=17), 3) triple-vCD tandem repeats (n=4), 4) vCD fusions to VP16^6^, and 5) vCD fusions to VP7 or VP28^58^. To each class we added mutational and partial sequence inversion variants to alter the vCD structure and gain mechanistic insights into vCD’s mode of activation.

**Fig. 3.**
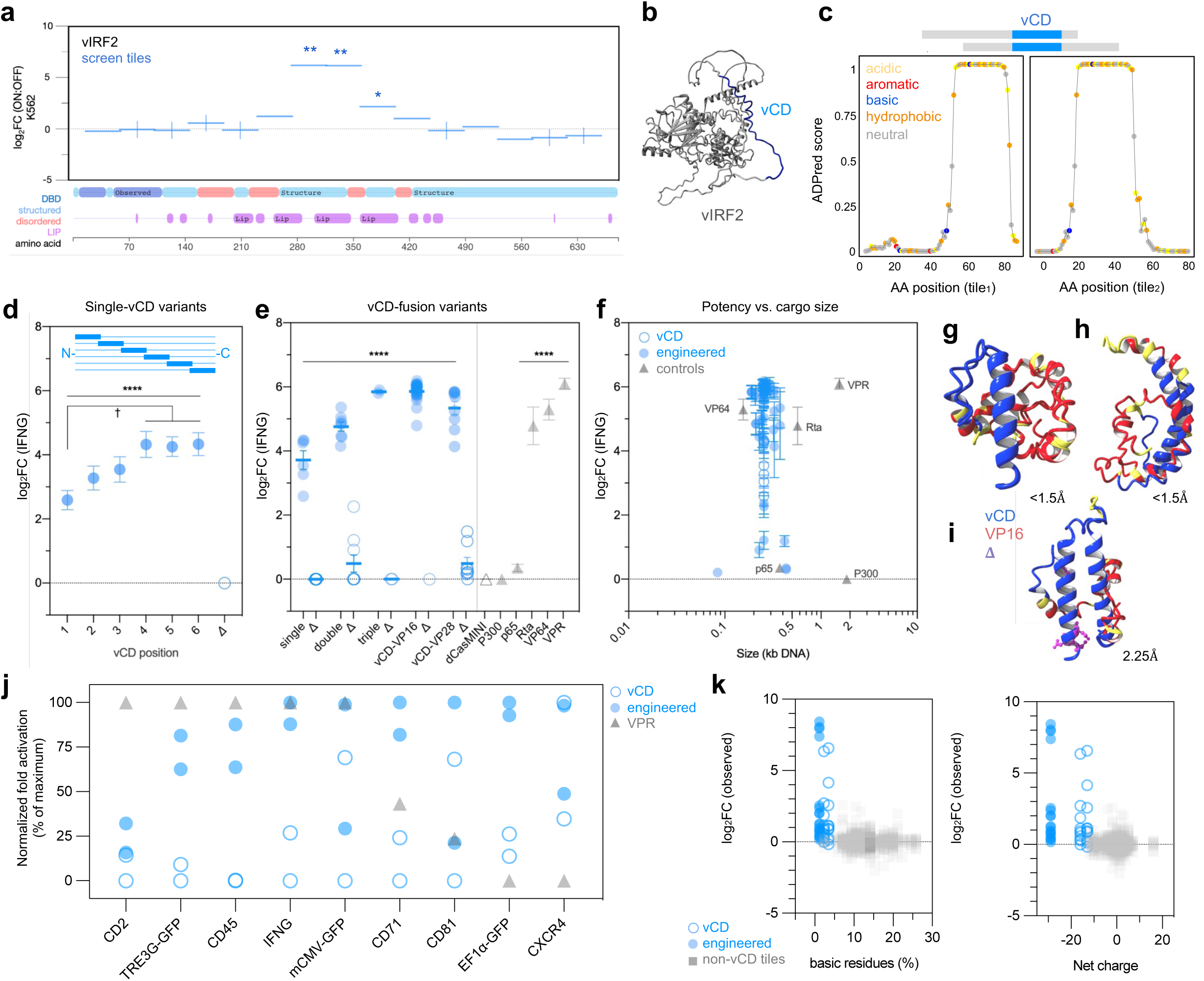
Engineered potency of vIRF2 core domain (vCD)-based modulators. **a,** Positional mapping of vIRF2 peptide tiles (x-axis) and respective activation strengths (log_2_ fold-change of ON:OFF barcode enrichment scores, mean ±SEM, **P<1.72×10^-61^, **P<1.72×10^-61^) in the initial mCMV-GFP K562 pooled screen (top). MobiDB schematic (below) indicates vIRF2 (Q2HR71) native protein features including structured (blue) versus disordered (red) regions, DNA-binding domain (DBD, dark blue), and linear interacting peptides (LIP, purple). **b,** 3-dimensional structure prediction of the full-length native vIRF2 and predicted vIRF2 core domain (vCD) (blue). **c,** Minimal vCD prediction by ADPred. The 32-a.a. vCD shared by the top two screened vIRF2 tiles, shown in (**a**), is enriched for acidic and hydrophobic residues. **d,e,** Activation potencies at IFNG (mean±SEM of 3 or more replicates per modulator; *P<0.001; †P<0.05, one-way ANOVA) of single-vCD modulators with the vCD at six positions of increasing distance from the dCasMINI and respective mutants (**d**), and multi-vCD fusions and respective mutants with benchmark activators (gray) (**e**). **f,** Activation potencies at IFNG (mean±SEM) of all engineered vCD-based modulators (n=101) (blue) and benchmarks (gray) relative to peptide coding size (DNA kb) (x-axis), with high-potency domains as compact as 64-a.a. **g,h,** 3-dimensional structure predictions of high-potency vCD modulators with close alignment of vCD cores (RMSD <1Å) contributing to form a stable vCD helix, with the postulated interface of charged residues exposed to the solute. **i,** 3-dimensional structure predictions of weak activator variants with mutations C4L, L5D, M7D, and L19D (RMSD 2.25Å). **j,** Activation potencies of two vCD-fusion variants at ten target contexts, relative to non-engineered vIRF2 tiles and VPR (gray), normalized to the maximum potency (expressed as %) observed in each experiment. **k,** Biochemical feature scores (x-axis) plotted against mean activation potencies at ten target contexts (y-axis) of vCD-fusions, non-engineered vCD tiles, and other non-vCD tiles.

We subjected the engineered vCD variant set (n=101) to arrayed activator testing at multiple genes in HEK293T cells and observed a wide range of potency effects. At each target, engineered vCD-based modulators (up to 98-a.a.) outperformed the unmodified vCD (85-a.a.) and, at six of nine targets, matched or outperformed VPR (523-a.a.) in potency (**Fig. 3j** and **Extended Data Figs. 4,5,7**). Potency increased with more negative electrostatic potential (ESP) but was dampened by amino acid substitutions in the most negative ESP variants (**Extended Data Fig. 7d**). We calculated a normalized robustness score based on activity at these targets, finding positive correlations with higher helix score and lower coil score (**Extended Data Fig. 7e**). Among single-core variants, stepwise increases in activation strength accompany the N-to-C distal shuffling of vCD, however substitutions of native vCD-flanking sequence resulted in total loss of activation strength (**Fig. 3d**). Duplicate and triplicate fusions led to stepwise increases in activation potency, and partial vCD inversions or single amino acid substitutions were largely tolerated, while multiple substitutions ablated activation (**Fig. 3e**). vCD-VP16 fusions and vCD-VP28 fusions gave the highest potencies, comparable to VPR, VP64, and Rta (**Fig. 3e**). Examination of 3-dimensional structure predictions revealed that the strongest activator fusions shared an exceptionally close alignment of vCD cores (RMSD<1.5Å) in a stable vCD ɑ-helix, with a postulated exposed interface of charged residues (**Fig. 3g,h**). In contrast, the weakening effects of multiple substitutions (RMSD=2.25 Å) suggests these residues collectively, though not singly, are critical to maintain helical or inter helix stability (**Fig. 3i**).

Analyzing amino acid composition, we find that vCD-VP64 fusions yield emergent activator-predictive properties in a super-additive manner beyond the ranges of either VP64 or single-vCD, namely in basic residue depletion, enrichment of acidic, turn percent, B-factor values, helix percentage, and a greater negative net charge (but not ESP) (**Fig. 3k, Extended Data Fig. 7f**). Taken together, we find that engineered vCD activators achieve functionally equivalent or greater potency and context-robustness compared to much larger benchmarks by enhancing the biochemical properties intrinsic to the unmodified vCD.

### Context-dependent durability of engineered activators

We next set out to evaluate the temporal activation kinetics of engineered variants relative to benchmarks (VPR, VP64, Rta, and P300) following transient transfection. We selected four human genes with diverse expression profiles and chromatin contexts: CD45, IFNG, CXCR4, and CD81 (**Extended Data Fig. 8a**) and performed time-series analyses in HEK293T cells with protein measurements by flow cytometry or ELISA as well as mRNA detection by RT-qPCR. Following transfection, we first confirmed that dCasMINI-modulator plasmid expression became undetectable between 6 and 9 days post-transfection (d.p.t.) (**Fig. 4a,b**).

**Fig. 4.**
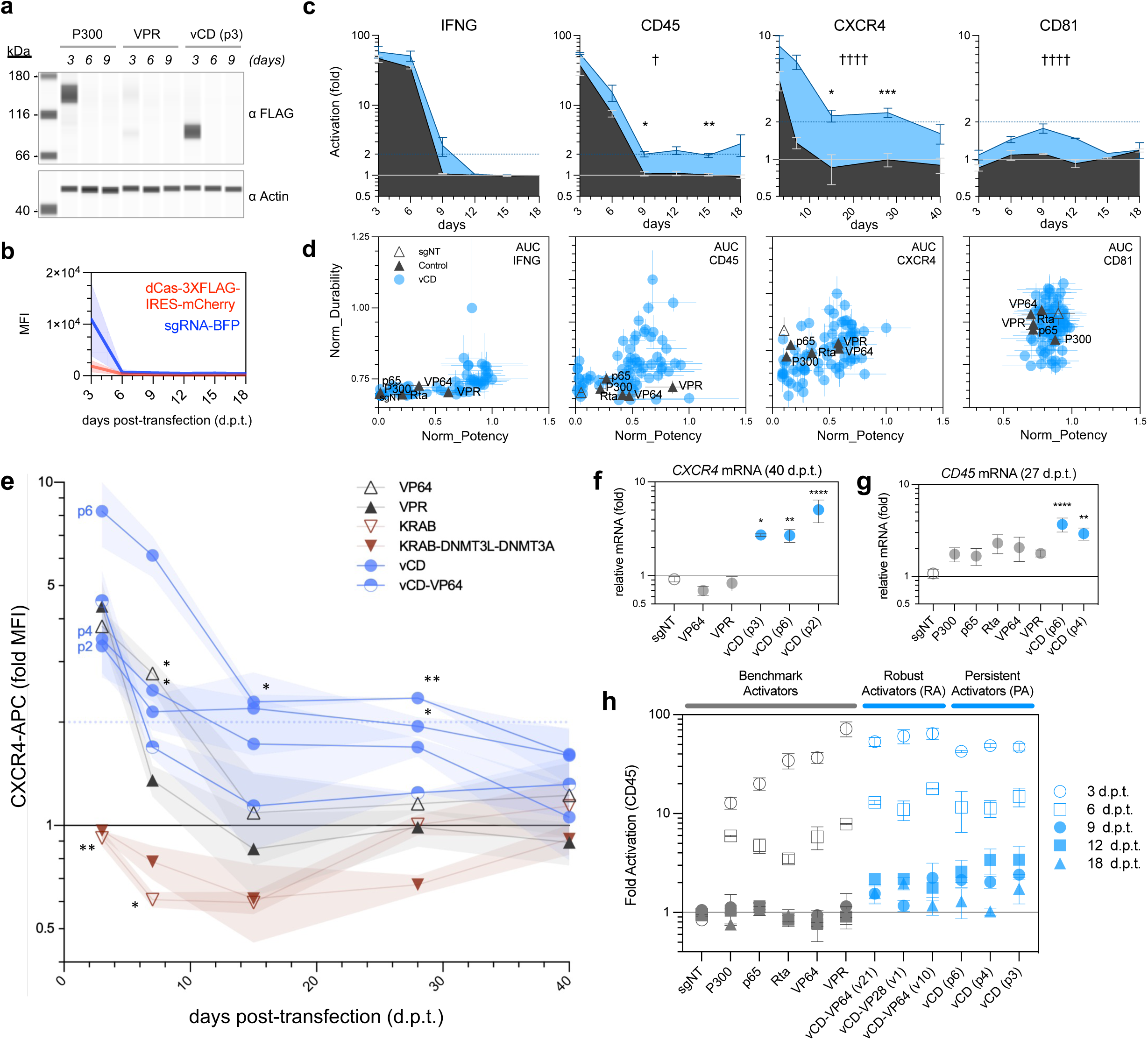
Context-dependent durability of engineered activators. **a,** Immunostaining to detect expression of 3XFLAG-tagged dCasMINI-modulator fusions following transient plasmid transfection in HEK293T cells at 3-9 d.p.t. **b,** Detection of plasmid loss by 6-d.p.t. by flow cytometry for dCas-modulator-IRES-mCherry and sgRNA-BFP plasmids. **c,** Target fold changes in activation (mean±SEM, n=3 or more per modulator) at each locus following transient recruitment of an individual vCD-based modulator versus VPR benchmark for the full time series: 3-to-29 d.p.t. for CXCR4 and 3- to 18-d.p.t. for IFNG, CD45, and CD81 (significance from VPR for the series, †P<0.05, ††††P<0.0001, and individual time points *P<0.05, **P<0.01, ***P<0.01, two-way ANOVA). Time series AUC indicated for normalized activator score calculations. **d,** Normalized durability scores (9 d.p.t. forward) at all four targets (±SD) (y-axis) versus normalized potency scores (3 and 6 d.p.t.) (±SD) (x-axis) for benchmarks VPR, VP64, Rta, p65, and P300 (black) and vCD modulators (n=101) (blue) with engineering classes indicated by shape. **e,** CXCR4 full time series comparing fold-change activations (CXCR4-APC geometric MFI, mean±SEM of 3 or more replicates, significance from VPR, *P<0.05, **P<0.01, two-way ANOVA) at each time point for single-vCD variants and controls. **f,** *CXCR4* mRNA detection (mean±SEM of 4 replicates, significance from sgNT, ****P<0.0001, **P<0.01, *P<0.05, one-way ANOVA) at 40 d.p.t. by indicated modulators following transient plasmid transfection in HEK293T cells. **g,** *CD45* mRNA detection (mean±SEM of 6 replicates, significance from sgNT, ***P<0.0001 *P<0.01, one-way ANOVA) at 27 d.p.t. by indicated modulators following transient plasmid transfection in HEK293T cells. **h,** CD45 activation data (normalized fraction of CD45-APC+ cells) at indicated time points for selected modulators (blue) and benchmarks (gray) following transient transfection. Early time points where modulator and sgRNA plasmids remain expressed (empty symbols) are distinguished from later time points with undetectable plasmid expression (filled symbols). vCD-fusion based robust activators (RA), vCD persistent activators (PA), and benchmark activators are indicated.

Benchmark activators VPR, VP64, and Rta consistently achieved high initial activation potencies at IFNG, CD45, and CXCR4 (3 d.p.t.), followed by acute drops in activity between 3 and 9 d.p.t., concurrent with loss of mCherry and 3xFLAG detection, down to baseline levels indistinguishable from negative controls (**Fig. 4c-e, Extended Data Fig. 8b-f**). Using this threshold, we calculated potency scores based on 3- and 6-d.p.t. values, and durability scores based on values at 9 d.p.t. and later values. In contrast to benchmarks, subsets of vCD-based modulators maintained durable activation (>2 SD above benchmarks) in a target-dependent manner (26.1% at IFNG, 43.5% at CD45, 44.6% at CXCR4, 17.4% at CD81) (**Fig. 4d**). At CD45 and CXCR4, we find that single-vCD variants can sustain a ∼2-fold activation plateau to 27 d.p.t. and 40 d.p.t., respectively (**Fig. 4e-h, Extended Data Figs. 8,9**). Importantly, we measured cell division rates via S-phase cell labeling and observed no significant changes (**Extended Data Fig. 8g**). We found that activation durability is not a direct function of initial potency. Persistent activators (PA) displayed moderate potency at early time points, but most consistently maintain target elevation more durably than high-potency robust activators (RA) (**Fig. 4h**).

Previous reports described a role for the KSHV vIRF2 protein in recruitment of CBP/P300 acetyltransferase complexes to target genes^56,59^. Reasoning that such a mechanism, if mitotically stable, might require “reading” of acetylation by bromodomains such as BRD4 and related BET family proteins^46,47,60^, we asked whether small molecule inhibitors (BRDi) might affect VCD-mediated gene activation. We performed a time series of CD45 activation, now with the added application of BRDi and other small molecule inhibitors (see **Methods**) against WDR5/MLL histone methyltransferase, EZH2/PRC2 demethylase, histone deacetylase, and DNA methyltransferase. In DMSO control conditions, we again observed that vCD activators durably sustained the highest CD45+ cell fractions (**Extended Data Fig. 9a,b**). In contrast, vCD-treated cells given BRDi drugs showed reduced CD45 activation (**Extended Data Fig. 9b,c, Supplementary Fig. 1**) and similar effects occurred at the CXCR4 locus (**Extended Data Fig. 9d**).

### Robust and persistent activation of therapeutically relevant genes in T-cells

Next, we sought to evaluate vCD-based candidate persistent activators (PA) and robust activators (RA) relative to benchmarks (**Fig. 5a**) in additional contexts of compelling biomedical interest to evaluate their potential in therapeutic applications. To this end, we tested strategies for upregulation of interleukins, toward enhancing anti-tumor response in immuno-oncology therapies^66-68^.

**Fig. 5.**
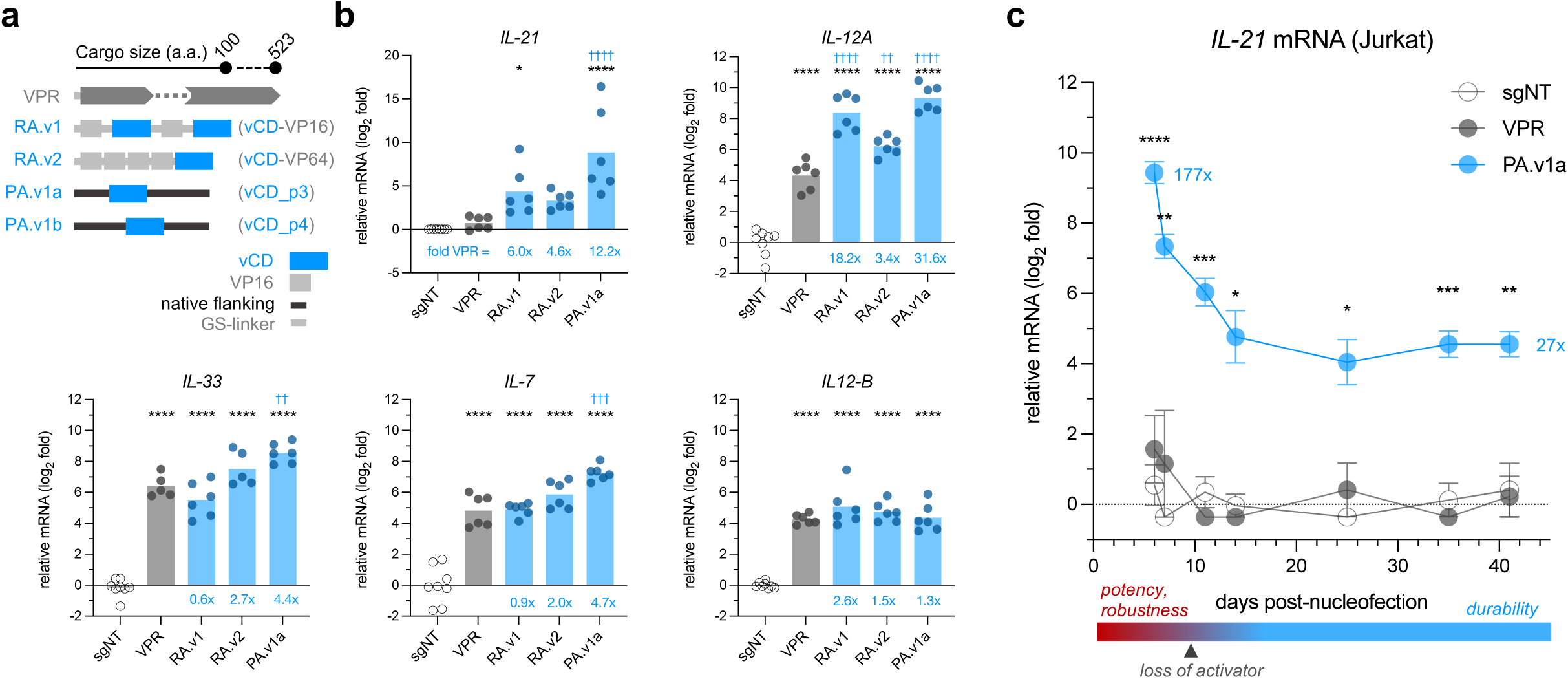
Robust and persistent activation of therapeutically relevant genes in T-cells. **a,** Schematic representations of vCD-fusion based robust activators (RA), vCD persistent activators (PA), and benchmark VPR. Boxes indicate the positions of vIRF2 core domain (blue) and VP16 domain (grey), and lines indicate vIRF2 native flanking sequence (black line) and GS-linkers (grey line). Relative cargo sizes (a.a.) are indicated relative to the benchmark activator VPR. **b,c** Target gene expression changes in dCas-modulator expressing Jurkat cells as measured by RT-qPCR, following sgRNA delivery by (**b**) lentiviral transduction at 7- and 10-days post-transduction (d.p.t.) or by (**c**) transient plasmid nucleofection with serial sampling across the indicated time series. Relative mRNA fold changes, normalized to non-targeting sgRNA conditions with the same modulators, are plotted as mean±SEM of 3-6 replicates. Significance from sgNT, ***P<0.0001, **P<0.001, *P<0.01. Significance from VPR, ††††P<0.0001, †††P<0.001, ††P<0.01, †P<0.05. One-way ANOVA (**b**) or two-way ANOVA (**c**). Folds (in blue) indicate fold versus VPR. **c,**bottom, Schematic representation of the brief potency and robustness window of benchmark activator VPR, and the sustained durability window of activation by a vCD persistent activator.

We selected seven interleukin genes whose upregulation would be clinically desirable in CAR-T cell armoring^71^, interleukin-21 (*IL-21*), interleukin-33 (*IL-33*), interleukin-12A (*IL-12A*), interleukin-12B (*IL-12B*), interleukin-7 (*IL-7*), interleukin-15 (*IL-15*), and interleukin-18 (*IL-18*), screened for high-efficiency sgRNAs for each target, and compared their activation by three lead vCD-based activators against that of the high-potency benchmark activator VPR in T-cell lymphocyte-derived Jurkat cells. To first evaluate their potency, we lentivirally co-delivered dCas-modulators and sgRNAs into Jurkat cells, enriched the transduced cells to purity by antibiotic selection, and measured relative mRNA fold inductions at 7- and 10-days post transduction (d.p.t.) (**Fig. 5b** and **Extended Data Fig. 10a,b**). At five of seven gene targets, one or more vCD activators outperformed VPR by factors up to ∼32-fold. Furthermore, despite constitutive expression by lentiviral delivery, between 7 and 10 d.p.t., we observed a drop in VPR efficacy (mean drop -32%) while vCD efficacies increased by 112%, 124%, and 158%, respectively, over the same period (**Extended Data Fig. 10b**). Next, we performed a transient single-dose administration by plasmid nucleofection and serially monitored *IL-21* mRNA induction from 6 to 41 days post-nucleofection (d.p.n.). Here, at 6 d.p.n. the vCD outperformed VPR by ∼177-fold and maintained significant activation to 41 d.p.n. of 27-fold, while VPR activation faded to non-targeting control levels by 11 d.p.n. (**Fig. 5c**). Digital PCR confirmed a rapid, time-dependent loss of the nucleofected plasmid throughout the time series (**Extended Data Fig. 10c**).

### Persistent LDLR activation in liver cells *in vivo*

Next, we evaluated a novel therapeutic approach for familial hypercholesterolemia via persistent upregulation of human LDLR, both *in vitro* and *in vivo*—a strategy particularly relevant as *LDLR* haploinsufficiency constitutes 85 to 90% of genetically confirmed cases^69,70^. We screened for high-efficiency *LDLR* targeting sgRNAs and performed time series analysis of LDLR activation following transient plasmid transfection, comparing vCD activators against the benchmarks VPR and VP64. vCD activators outperformed VP64 by ∼75% and ∼90% at 2 and 16 d.p.t., respectively, achieving LDLR surface protein levels comparable to those induced by a positive control *LDLR* overexpression cDNA plasmid (**Fig. 6a,b**). In a long-term experiment, we compared activation by vCD and VPR, finding that vCD and VPR resulted in *LDLR* mRNA levels of 2.1-fold and 1.3-fold, respectively, at 70 d.p.t. (**Fig. 6c**).

**Fig. 6.**
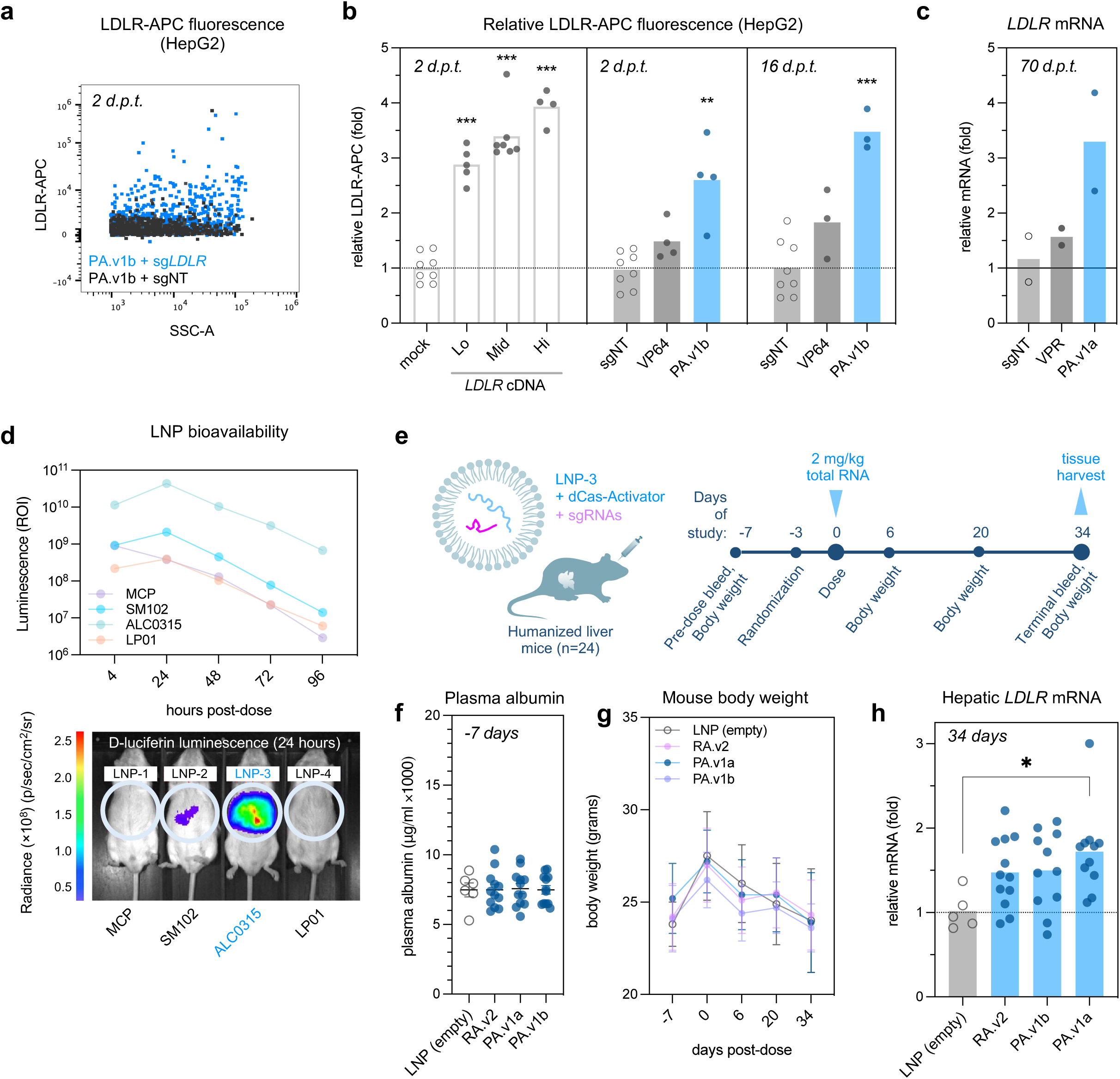
Persistent activation *in vitro* and *in vivo*. **a-c,** LDLR activation in HepG2 cells following transient plasmid transfection. Representative flow cytometry dot plot (**a**) comparing LDLR-APC surface protein fluorescence in dCas-modulator expressing HepG2 cells treated with LDLR-targeting (blue) or non-targeting control (gray) sgRNAs, and quantification of changes in LDLR-APC mean fluorescence intensity at the indicated time points following transient transfection (**b**) (mean±SEM of 3-8 replicates; significance calculated against non-targeting controls, ***P<0.001, **P<0.01, *P<0.05, one-way ANOVA). cDNA encoding the LDLR open reading frame was separately transfected in untreated HepG2 cells to compare LDLR-APC fluorescence levels in a classical gene over-expression context. **c,** *LDLR* mRNA activation in HepG2 cells measured by RT-qPCR at 70 days following transient co-transfection of dCas-modulator plasmid and sgRNA plasmids. **d,** Quantified bioluminescence in mouse livers across the LNP selection experiment from 4 to 96 hours post-administration. Humanized mice were dosed with four different LNP formulations encapsulating D-Luciferase mRNA (n=1/LNP) and imaged for luminescence intensity from 4 to 96 hours using the InVivo Imaging System (IVIS). **e,** Study schema for *in vivo LDLR* activation experiment. Humanized liver mice (n=24) were bled and weighed at day -7, randomized at day -3 (n=6 mice per experimental group), and dosed by retro-orbital injection at day 0 with 2 mg/kg total RNA (mRNA+sgRNA, 1:1 wt/wt ratio) encapsulated into lipid nanoparticles (LNP). Intermediate bleeds were taken at indicated time points followed by terminal tissue harvest at day 34. **f,** Pre-randomization plasma albumin levels indicating uniform liver humanization levels across mice at day -7 pre-dose. **g,** Mouse body weight measurements obtained across the study time series. **h,** Hepatic *LDLR* gene expression was measured by RT-qPCR as fold change after normalizing to *RPL19* from humanized mouse livers at 34 days post LNP-administration (mean±SEM of 6-12 mice; significance calculated against LNP (empty), **P<0.05, *P<0.01, one-way ANOVA).

Non-viral therapeutic vectors such as lipid nanoparticles (LNP) offer improved safety profiles over AAV vectors^73^ and in recent years have been widely used to deliver synthetic mRNA cargos even in healthy individuals, but LNP-mediated dCas activation approaches will require that the therapeutic payload be capable of inducing long-lasting gene modulation past a transient single-dose administration. Previous *in vivo* studies in mice using LNP-delivered dCas-VPR and dCas-P300 targeted to liver genes have shown significant activation levels up to 3 days, with effects dissipating to baseline by 9 days^39^. We therefore set out to determine if the superior potency and durability profiles shown by vCD *in vitro* would be recapitulated *in vivo* via LNP-mediated single-dose administration in a humanized liver mouse model (**Extended Data Fig. 10d**). After empirically selecting an optimal LNP formulation for hepatic expression (**Fig. 6d**), we encapsulated synthetic mRNAs encoding dCas-vCD variants together with sgRNAs targeting human *LDLR* (**Fig. 6e**). Prior to dosing, mice were randomized according to body weight and human albumin levels as proxy for liver humanization (**Fig. 6f**). Mice were dosed via retro-orbital injection of LNPs with or without dCas-vCD mRNA and sgRNA cargos and subjected to serial blood sampling before a terminal tissue harvest at 34 days post-dose for extraction of hepatic mRNA (**Fig. 6e**). Digital PCR of hepatic mRNA confirmed that LNP cargos were below the level of quantitation. Clinical observations throughout the study showed no significant changes in serum alanine aminotransferase (ALT) or mouse body weights when compared to the randomized control group (**Fig. 6g** and **Extended Data Fig. 10e**), indicating that the therapeutic LNPs did not cause adverse changes in treated mice. We measured human *LDLR* expression in hepatic mRNA harvested at 34 days post-dose and found that, compared to the empty LNP condition, PA.v1a significantly activated *LDLR* levels by ∼72% (**Fig. 6h**), while PA.v1b and RA.v2 activated *LDLR* levels by ∼49% and 47%, respectively. Thus, in contrast to previous reports of temporally restricted activation by VPR under comparable conditions^39^, the compact (85 a.a.) vCD-based modulator PA.v1a, following a transient, single-dose LNP administration, was able to maintain human *LDLR* gene activation for a period up to five weeks.

## Discussion

A recent explosion in high-throughput screening studies has identified and characterized domains capable of eliciting desired changes in gene expression^17,18,19,27^. Typically, these studies have constrained the search space for such activities to known regulatory domains, screening with reporter genes at synthetically controlled but uncharacterized epigenomic contexts. This approach, while productive, risks limiting the utility of such domains as it does not account for the heterogeneity of chromatin contexts among the hundreds or thousands of disease-relevant target genes in human cells. By contrast, in this study we performed the first direct comparison of transcriptional regulators among tens of thousands of peptides sourced from divergent taxonomic lineages of extremophilic and mesophilic environments. We uncovered a marked activator potency bias for viral peptides and suppressor potency biases for human peptides and for thermophilic and acidophilic archaeal peptides. Amongst these potent viral activators, we discovered a sub-domain capable of extremely durable target gene activation, lasting several weeks post-transient delivery. We evaluated this activity in multiple cell types at endogenous human genes *in vitro*, and showed that it out-performed such well-characterized activators as VP64 and VPR in both potency and durability. The vCDs reported in this study exhibit robust activation of target genes, particularly those associated with quiescent or bivalent chromatin states (as indicated by promoter ChromHMM annotation) in Jurkat cells. This contrasts sharply with the performance of VPR at the same gene targets. Our study was designed to highlight the efficacy of compact persistent activators, successfully maintaining the activation of an endogenous target gene for five weeks in primary human hepatocytes *in vivo* following single dose via LNP delivery. By overcoming challenges observed in prior *in vivo* gene activation studies, including limited durability, cargo size constraints necessitating multiple AAVs for delivery in some cases, and the requirement for repeat administration of LNPs, we have effectively addressed these limitations. This work not only showcases the potential of our approach but also provides a practical roadmap for achieving therapeutic persistent gene activation.

We have shown that the organismal provenance of isolated peptide fragments confers significant differences in transcription modulatory function as defined by 1) potency, 2) chromatin context-independent robustness, and 3) durability of activity. Furthermore, our analysis of amino acid composition suggests distinct biochemical strategies by which different taxa have evolved unique transcriptional capabilities, forming the basis for predictive activator selection criteria previously unappreciated in the field. Using machine learning and insights from structural predictions, we identified biochemical and sequence-level peptide features of activation allowing us to develop a rational engineering strategy that enhanced the potency and durability of transcriptional activation. Together, these predictive insights may facilitate future directed screening efforts for the accelerated discovery and *de novo* design of epigenetic engineering tools^62^.

In biomedically-relevant contexts, variants of the vIRF2 core domain (vCD) discovered herein outperformed current CRISPR activation tools of far greater cargo size in transcriptional potency and demonstrated a unique ability to persistently activate genes *in vitro* and *in vivo*. Future mechanistic characterizations underlying this effect might test the hypothesis that the vCD acts as an “initiator” akin to the endogenous transcription factors that establish cellular identity, modifying the chromatin structure to confer stable transmission of gene expression programs. When applied in human cell contexts, viral activators may circumvent the obstacle of transcriptional cofactor scarcity and negate the rate-limiting competition for co-activating proteins in human cells^61^, thereby enabling the superior potency of vCD to larger multipartite fusions observed in multiple contexts. Our further demonstrations of mitotically stable gene activation, together with the practical advantages in deliverability of hypercompact dCas-vCD fusions, marks an important advance toward the realization of epigenetic editing as a therapeutic modality via persistent activation of disease-modifying genes *in vivo*.

---

**Extended Data Fig. 1.**
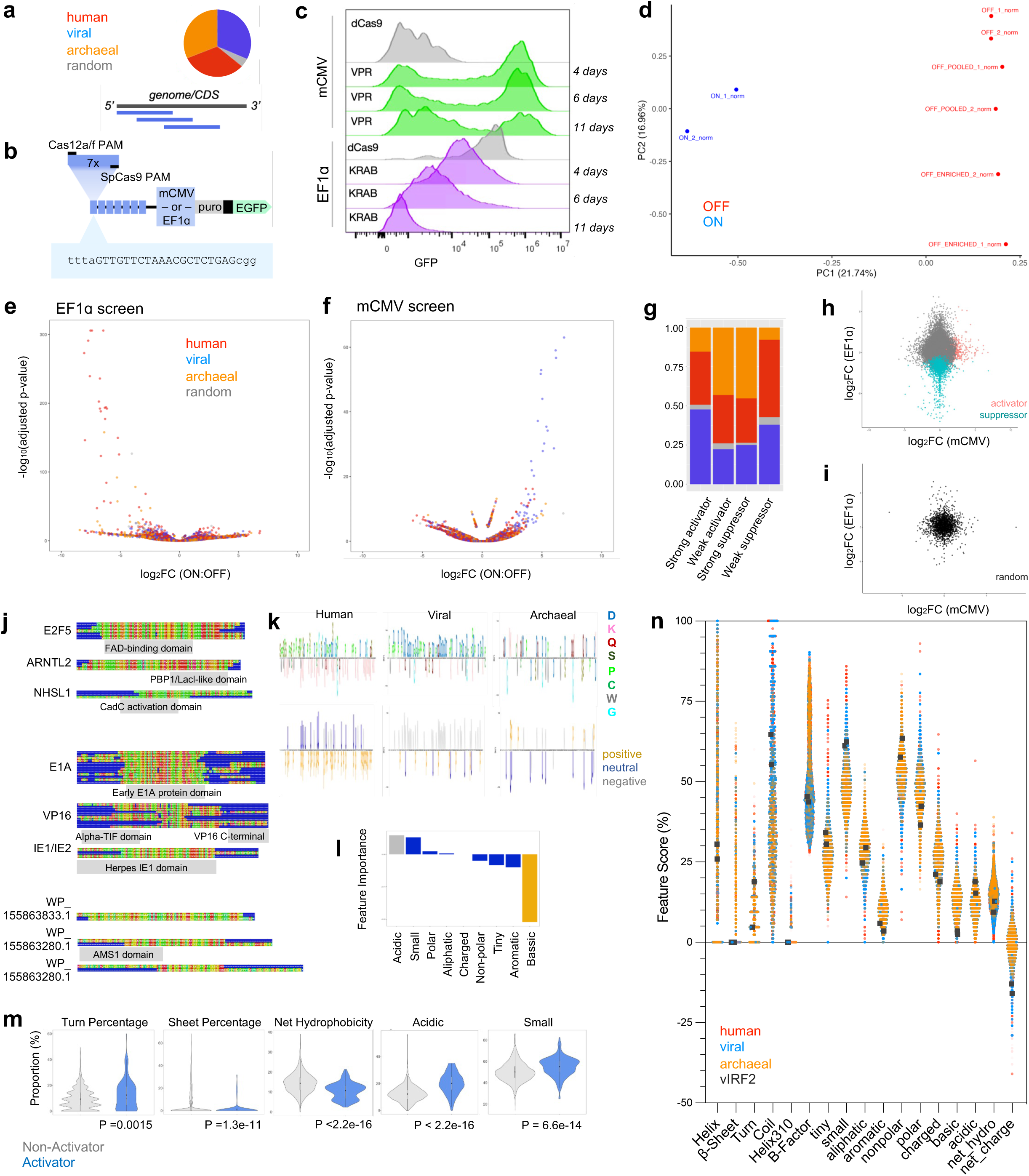
Pooled screening for human, viral, and archaeal modulators. **a,** Illustrations of initial high-throughput library composition and tiling strategy. 85-a.a. fragments were encoded as 300-mer DNA oligonucleotides, each labeled by a unique 12-mer DNA barcode. 549 human nuclear proteins were tiled with 50% sequence overlap to generate 16,139 oligos. Viral full genomes, human viral transcriptional regulators (hVTRs)^26^, metagenomic viral peptide predictions, and the archaeon full genome were tiled at 66.6% overlap to generate 27,799 oligos. GC content-matched scrambled sequences were also included as random negative controls. **b,** Schematic of custom transcriptional reporter constructs for screening activators (mCMV promoter with default GFP-OFF activity) and suppressors (EF1ɑ promoter with default GFP-ON activity). Each reporter construct contains seven identical sgRNA landing sites with dual-PAM sites for targeting via dCas9 or dCasMINI. **c,** Reporter validations in pilot studies monitoring flow cytometric GFP fluorescence in mCMV-GFP and EF1ɑ-GFP K562 cells bearing lentiviral integration of sgRNA-BFP vectors, sampled at indicated days post-lentiviral transduction of dCas9-VPR, dCas9-KRAB, or dCas9 without modulator fusion. **d,** Principal component analysis (PCA) of barcodes detected in pooled screen samples with separation of ON (n=2) and OFF (n=6) samples. **e,f,** Volcano plots of screen hits in EF1ɑ-GFP suppression screens (**e**) and mCMV-GFP activation screens (**f**), colored to illustrate library sources as human (red), viral (blue), archaeal (orange), or random (gray). **g,** Distributions of human, viral, archaeal, and random origins of strong/weak activators/suppressors defined by the highest and lowest quartiles of activators/suppressors respectively. **h,i,** Transcriptional effects (fold-change in ON:OFF barcode enrichment) of each modulator tile in both mCMV-GFP (x-axis) and EF1ɑ-GFP (y-axis) screens. Colors distinguish robust activators and suppressors (**h**), or random negative control tiles (**i**). **j,** Selected sequence alignments for human (top), viral (middle), and archaeal (bottom) sequence homology clusters illustrating common functional domains within activator clusters. **k,** Enrichment of amino acid residues composing human (n=209), viral (n=177), and archaeal (n=154) activators colored by residue type (top) and charge (bottom) calculated using Fisher’s exact test. **l,** Extracted feature importance from top generalized linear regression model detailing which residue types were predictive of activation strength (gray: acidic, gold: basic, blue: other). **m,** Violin plots comparing turn percentage, sheet percentage, net hydrophobicity, acidic residues, and small residues in activation hits (blue) and non-activator peptides (gray). P-values reported are based on Wilcoxon rank-sum testing. **n,** Biochemical feature distributions among screened human (red), viral (blue), and archaeal (orange) peptides. vIRF2 tiles (black square) are indicated.

**Extended Data Fig. 2.**
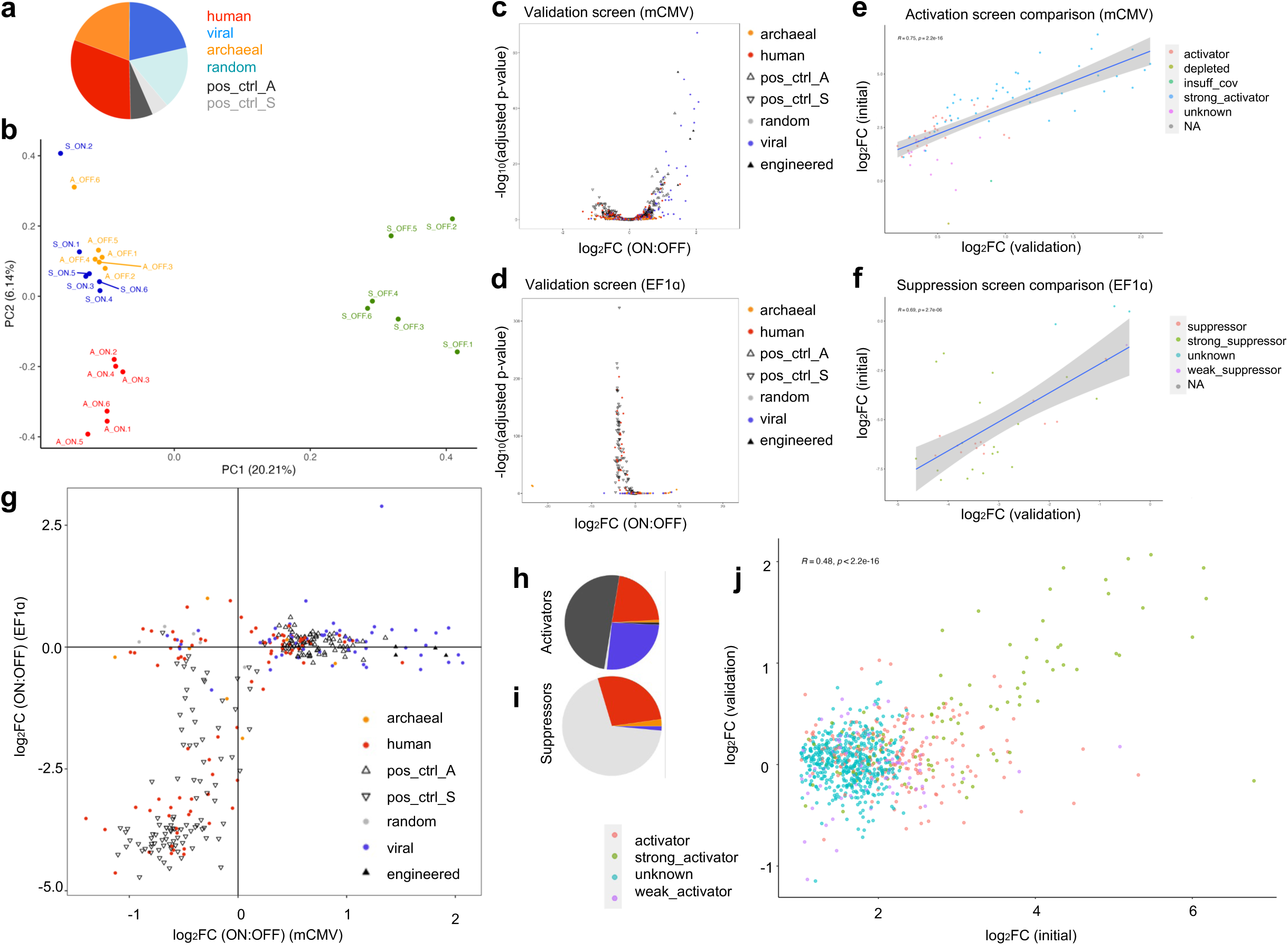
Modulator validation by pooled sub-library validation screening. **a,** Sub-library composition for the validation screens of putative modulators from the full-library screen (954 activators, 1,228 suppressors, and 22 dual-activity tiles) and selected engineered activators containing vIRF2-VP16 fusions. 949 negative controls included scrambled sequence tiles, tiles depleted in both full-library mCMV-GFP and EF1ɑ-GFP screens, and tiles initiated with stop codons. Positive control tiles were taken from published data^10,18^. **b,** PCA of modulator validation libraries. 6 replicate samples were analyzed for each of four conditions: mCMV-GFP-ON, mCMV-GFP-OFF, EF1ɑ-GFP-ON, and EF1ɑ-GFP-OFF. **c,d,** Volcano plots indicate screen hits for mCMV-GFP (**c**) and EF1ɑ-GFP (**d**) screens. Colors differentiate tile sources. Shapes differentiate test tiles (circle) from published positive controls (clear triangle) or engineered vIRF2-VP16 fusions (filled triangle). **e,f,** Correlation of screen hits detected in sub-library validation (x-axis) and full-library (y-axis) for both mCMV-GFP (**e**) and EF1ɑ-GFP (**f**) screens. **g,** Context robustness of validation screen hits at both mCMV-GFP and EF1ɑ-GFP promoters. Dots are colored by taxonomic origin (red: human, blue: viral, orange: archaeal, gray: random) and positive controls for activators and suppressors are represented by triangles and inverted triangles, respectively. **h,i,** Distributions of tile provenance of validated mCMV-GFP activators (**h**) and EF1ɑ-GFP suppressors (**i**). **j,** Correlation of mCMV-GFP activators in full-library (x-axis) and sub-library validation (y-axis), colored by classification as strong or weak activators by barcode enrichment.

**Extended Data Fig. 3.**
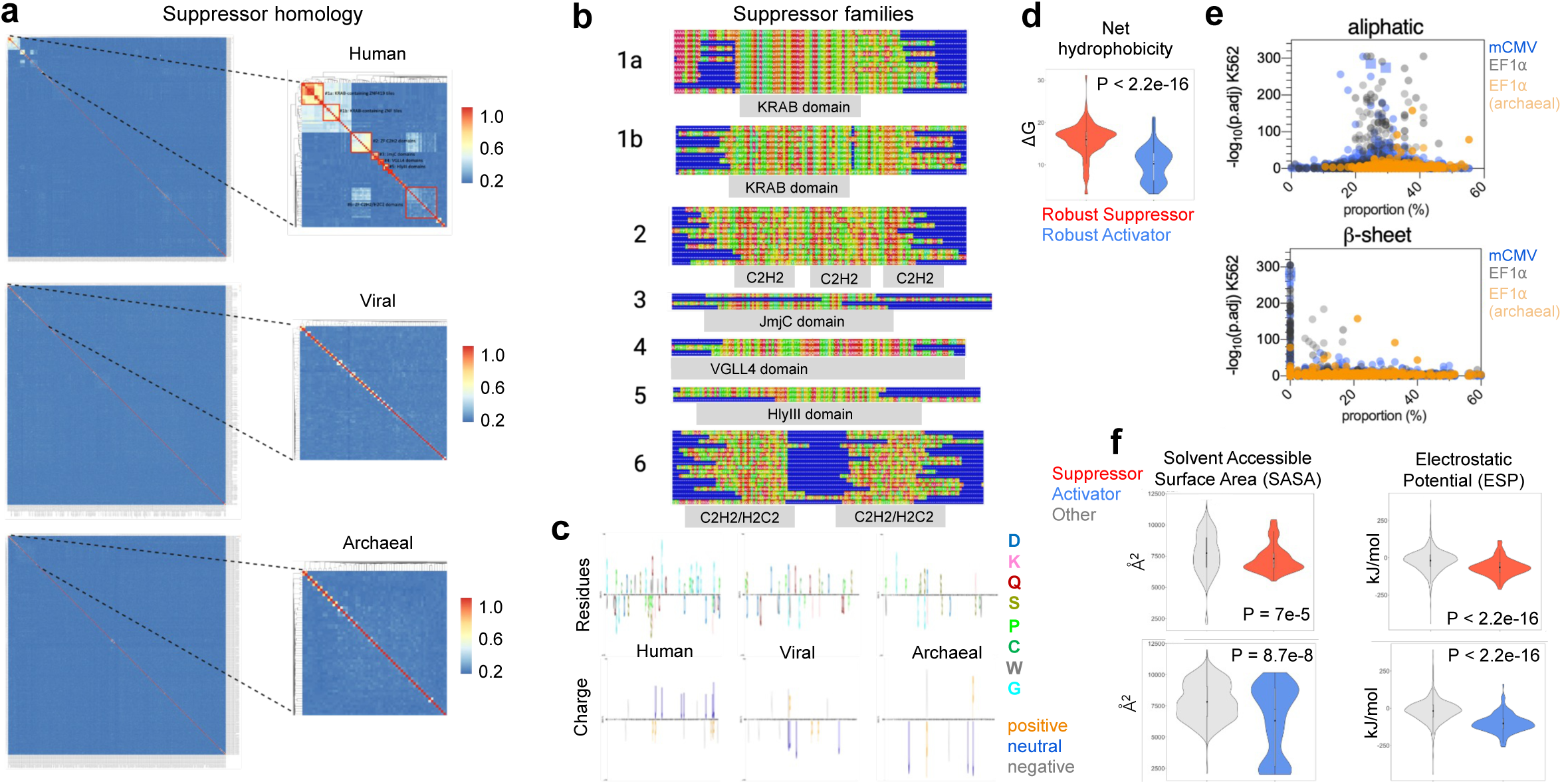
Clustering, alignment, and biochemical enrichment of suppression tiles. **a,** Sequence homology clustering of human (top), viral (middle), and archaeal (bottom) suppressors identified in the EF1ɑ-GFP high-throughput screen. **b,** Selected sequence alignments for human (top), viral (middle), and archaeal (bottom) sequence homology clusters illustrating common functional domains within suppressor clusters. **c,** Enrichment of amino acid residues composing human, viral, and archaeal suppressors colored by residue type (top) and charge (bottom) calculated using Fisher’s exact test. **d,** Violin plot comparing net hydrophobicity in robust activators (blue) and robust suppressors (red). P-values reported are based on Wilcoxon rank-sum testing. **e,** Biochemical feature scores (x-axis) plotted against hit significance (ON:OFF, adj. p-val) for the full modulator libraries in mCMV-GFP and EF1ɑ-GFP K562 screens. Scores in mCMV-GFP (blue) and EF1ɑ-GFP (gray). vIRF2 activator hits at mCMV-GFP (triangles), and archaeal suppressor hits at EF1ɑ-GFP (orange), are indicated. **f,** Violin plots comparing solvent accessible surface area (SASA) and electrostatic potential (ESP) in suppressors (red) vs. non-suppressors (gray) (top) and activators (blue) vs. non-activators (gray) (bottom). P-values reported are based on Wilcoxon rank-sum testing.

**Extended Data Fig. 4.**
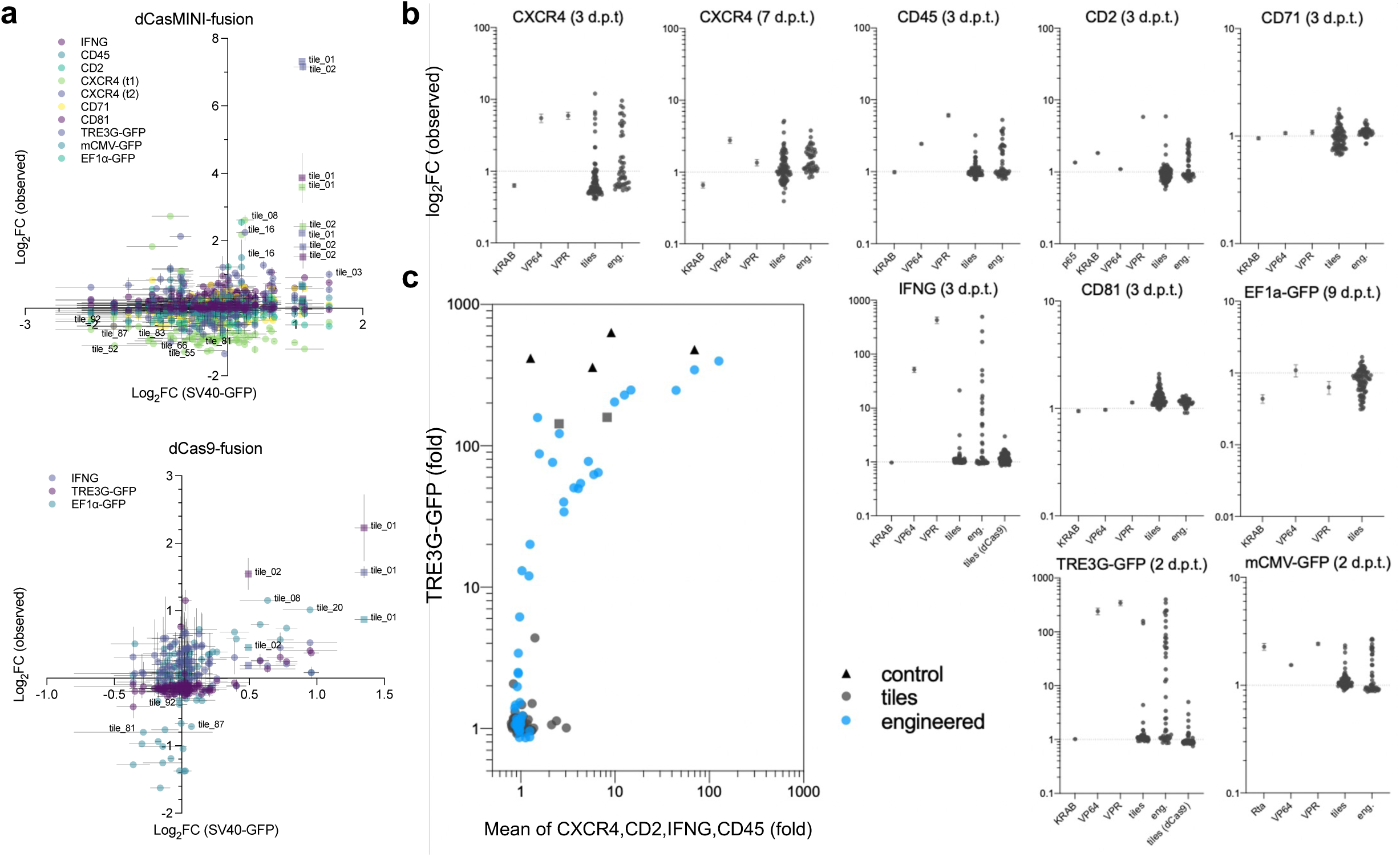
Modulation of protein expression in individual recruitment assays. **a-c,** Observed transcriptional effects on protein expression in HEK293T cell experiments (summarized in Fig. 2) as measured by flow cytometry or ELISA. Selected viral and archaeal modulator screen tiles (n=95) were targeted to synthetic and endogenous gene promoters via dCasMINI (**a,**top) or dCas9 (**a,**bottom) recruitment. Plots compare modulation of SV40-GFP (x-axis) against modulation at indicated targets (y-axis). Dots represent observed modulator activity per target (mean±SEM of 3 or more replicates) in observed protein expression, relative to non-targeting sgRNA (sgNT) and dCasMINI recruited without modulator fusion (dCasMINI) conditions. **b,** Individual experiments at various targets shown in (**a**) testing viral and archaeal modulator screen tiles (n=95), benchmark modulators VPR, VP64, Rta, p65, KRAB, and engineered variants based on viral activator tiles (tile_1 and tile_2) from vIRF2. **c,** Correlation of TRE3G-GFP activation against the mean activation of four endogenous genes indicated, per modulator.

**Extended Data Fig. 5.**
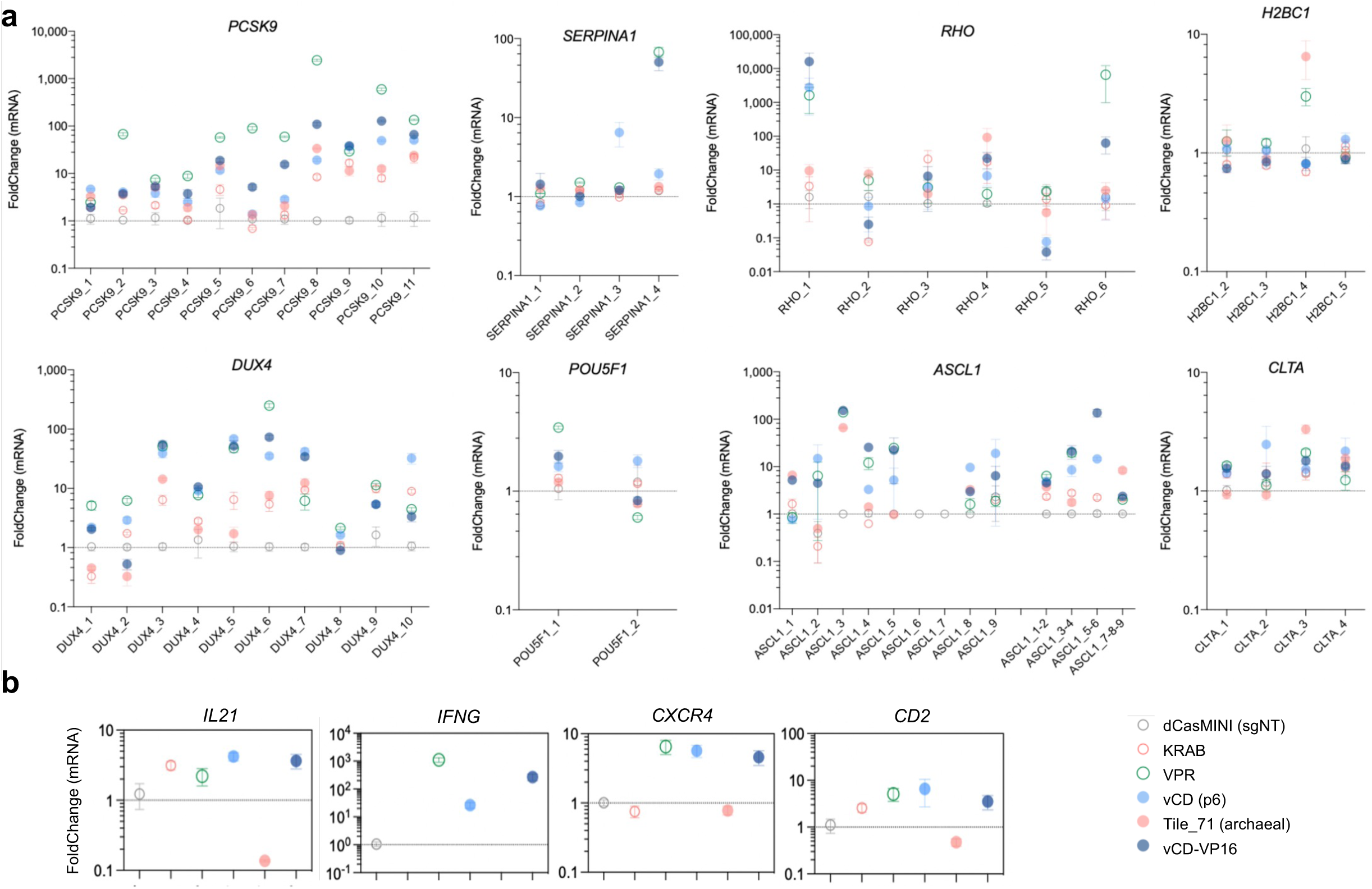
Modulation of mRNA expression in individual recruitment assays. **a,b,** RT-qPCR results (fold-change mean±SEM) at indicated genes HEK293T cells sampled at 3 d.p.t. following plasmid transfections (n=4) of dCasMINI without modulator (empty circle), or dCasMINI fusions to KRAB (red, empty), VPR (green, empty), vIRF2 screen tile (light blue), archaeal suppressor tile_71 (red), or engineered vCD-VP16 fusion (dark blue). dCasMINI-modulator plasmids were co-transfected with either individual sgRNA plasmids (**a**) or multiplexed sgRNA plasmid pools (**b**).

**Extended Data Fig. 6.**
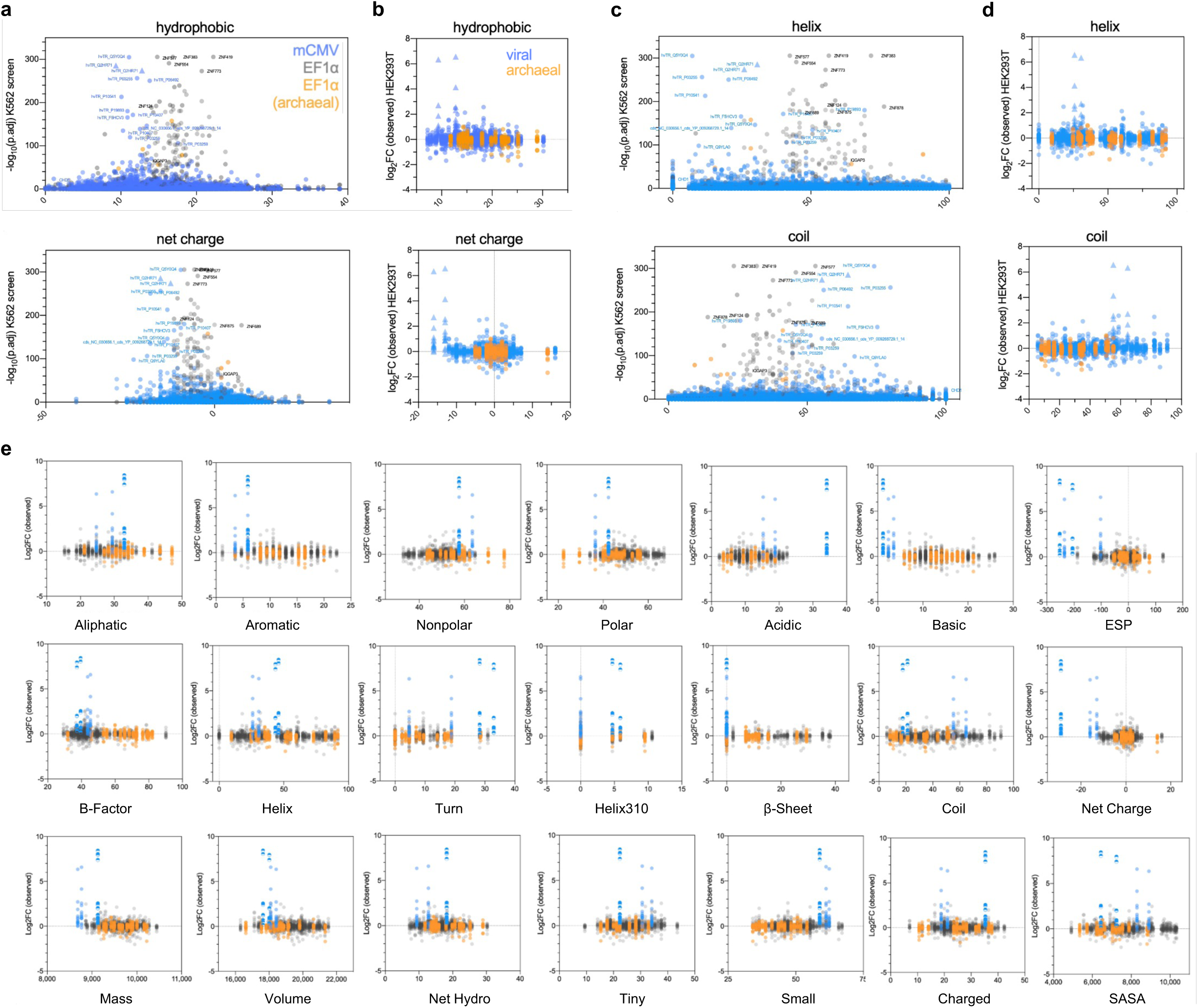
Biochemical feature correlations with robust modulator performance. **a-d,** Side-by-side comparison of K562 screen predictions for feature importance on modulator activity (Fig. 1c,d) and observed functional testing in HEK293T cells. (**a,c**) Biochemical feature scores (x-axis) plotted against hit significance (ON:OFF, adj. p-val) for the full modulator libraries in mCMV-GFP and EF1ɑ-GFP K562 screens. Scores in mCMV-GFP (blue) and EF1ɑ-GFP (gray). vIRF2 activator hits at mCMV-GFP (triangles), and archaeal suppressor hits at EF1ɑ-GFP (orange), are indicated. (**b,d**) Indicated biochemical feature scores (x-axis) are plotted against observed protein activation fold-changes (mean of technical triplicates) for 95 selected viral (blue) and archaeal (orange) modulators in ten experiments of dCasMINI-modulator fusions co-transfected in HEK293T cells with targeting sgRNAs for either IFNG, CD45, CXCR4, CD2, CD71, CD81, TRE3G-GFP, SV40-GFP, mCMV-GFP, or EF1ɑ-GFP. vIRF2 activator hits at mCMV-GFP are indicated (triangles). **e,** Biochemical feature differences between vIRF2 activator tiles and engineered vCD-VP16 fusions. Feature scores (x-axis) are plotted against observed protein activation fold-changes, adding the activity and scores of two engineered vCD-VP16 fusions (blue semi-circles) and highlighting the original vIRF2 screen tiles (blue). Archaeal suppressor hits (orange) are indicated.

**Extended Data Fig. 7.**
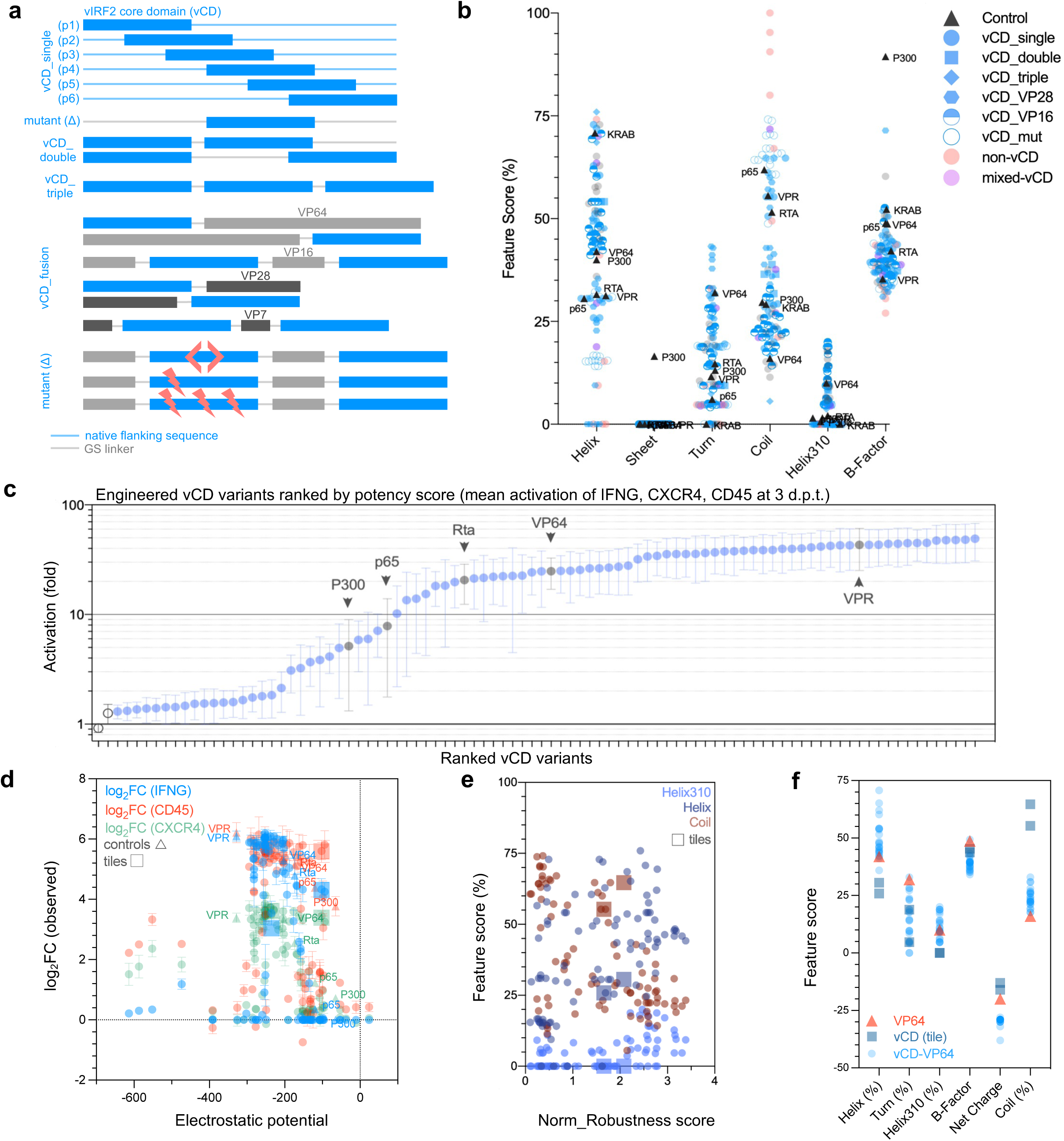
Domain-based engineering of hypercompact activators. **a,** Schematics illustrate engineered variants containing the 32-a.a. vCD, testing various configurations of N-to-C position, fusion partners, linker sequences, vCD inversions, and vCD mutations. **b,** Biochemical properties altered by engineering the vCD domain. **c,** Summarized activation potencies following co-transfection in HEK293T cells (n=3 or more per modulator) with targeting sgRNA plasmids and dCasMINI fusions to all engineered vCD variants (n=101) relative to positive controls VPR, VP64, Rta, p65, and P300. dCasMINI without modulator and paired with targeting sgRNAs, and dCasMINI-VPR paired with non-targeting sgRNA served as negative controls for normalization and fold-change calculations. Y-axis values are an averaged protein activation score for three experiments targeting CD45, CXCR4, and IFNG (mean±SEM). **d,** Activation potencies (mean±SEM of 3 or more replicates per modulator) of engineered vCD-based modulators (n=101) at IFNG, CD45, and CXCR4 at 3 d.p.t., relative to electrostatic potential of each modulator. **e,** Normalized robustness score at IFNG, CD45, and CXCR4 (y-axis), relative to percent residue enrichments for helix and coil structures. **f,** Changes in biochemical feature scores of vCD-VP64 fusions relative to scores of either VP64 or single-vCD modulators alone.

**Extended Data Fig. 8.**
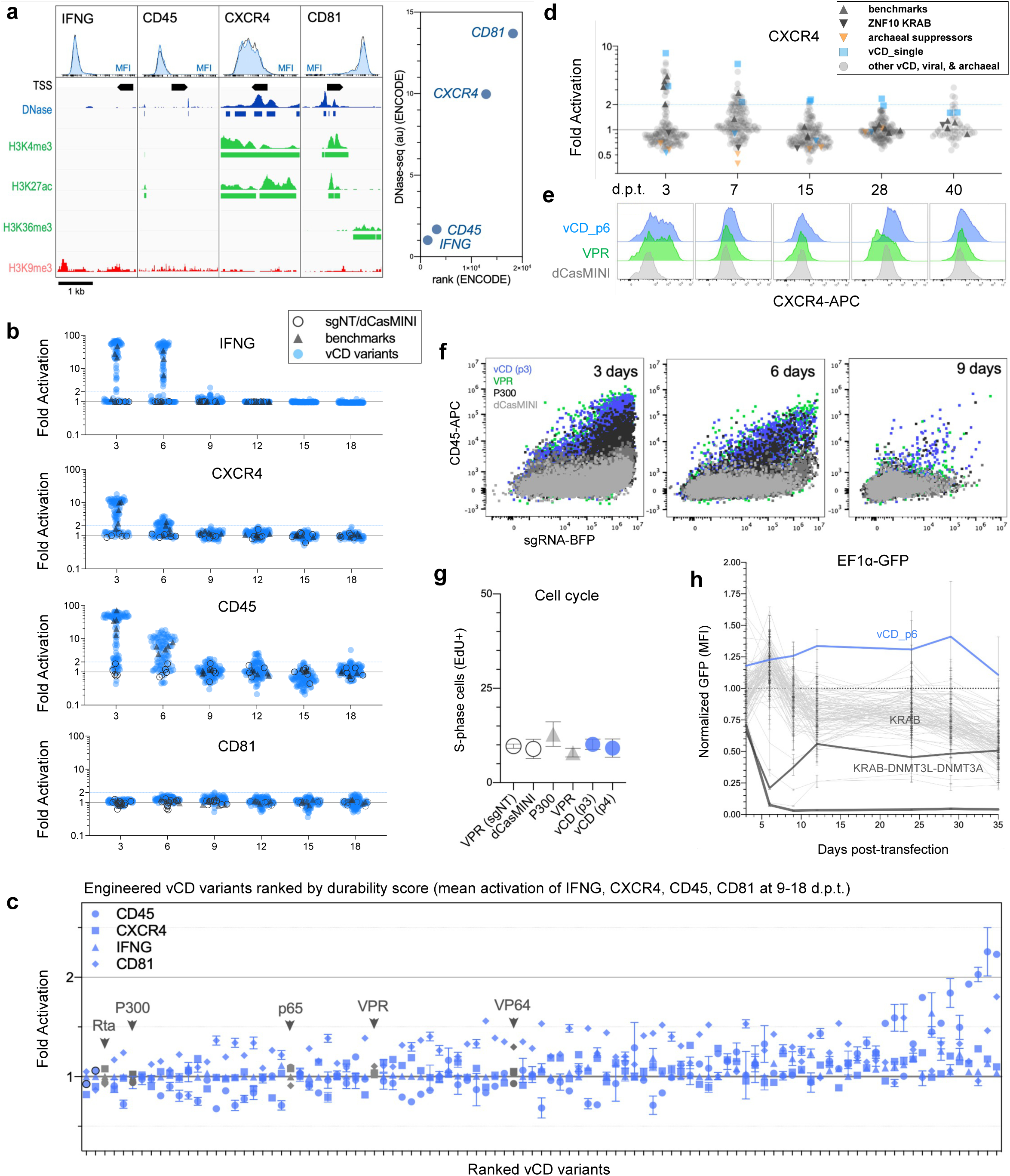
Prolonged activation kinetics of hypercompact activators. **a,** Chromatin contexts and HEK293T flow cytometric baselines of surface protein expression (left) and ENCODE DNAse-seq rankings (right) for CD45, IFNG, CXCR4, and CD81 genes targeted for activation time series measurements. **b,** Arrayed durability screening in HEK293T cells following transient transfection of dCasMINI-modulator fusions and targeting sgRNA plasmids. Observed fold-changes per modulator in secreted protein by ELISA (IFNG), APC MFI (CXCR4, CD81), and normalized fraction of CD45-APC+ cells (CD45) in HEK293T cells at indicated time points (x-axis). Dots represent means per modulator. Positive controls (VPR, VP64, Rta, p65, P300) were transfected in 5-10 replicates. Normalization (negative) controls (n=29 replicates per experiment) were dCasMINI without modulator fusion with (n=5) and without (n=2) targeting sgRNAs (sgT), and non-targeting sgRNA (sgNT) with dCasMINI-modulator fusions to VPR (n=4), VP64 (n=2), Rta (n=2), p65 (n=2), P300 (n=2), vCD_single (n=4), vCD-VP16 (n=4), and vCD-VP64 (n=2). Test modulators were vCD-based variants (n=101) transfected in 2-3 replicates. **c,** Summarized durability per target for all modulators at post-plasmid time points (mean±SEM of fold changes from 9-18 d.p.t.) at IFNG (triangle), CD45 (circle), CXCR4 (square), and CD81 (diamond). Ranked by the total mean of values across four targets, i.e. contextual robustness and durability. Benchmarks indicated (gray, arrows). **d,e,** Independently repeated CXCR4 experiment testing engineered modulators (n=48), viral and archaeal screen tiles (n=95), and benchmarks. Observed mean fold-changes (**d**) in CXCR4-APC shown per modulator with representative CXCR4 flow cytometry data (**e**) at 3-40 d.p.t. for VPR (green), vCD_p6 (blue), dCasMINI without modulator fusion and targeting sgRNAs (gray). **f,** Representative CD45 flow cytometry data at 9 d.p.t. for VPR (green), P300 (black), vCD_p3 (blue), dCasMINI without modulator fusion and targeting sgRNAs (dark gray), and VPR with non-targeting sgRNA (light gray). **g,** Cell cycle analysis of S-phase labeled cells following pulse-chase of EdU and flow cytometric EdU detection at 8 d.p.t. of indicated modulator plasmids. **h,** Observed fold-changes in EF1ɑ-GFP fluorescence in HEK293T cells (mean±SEM) measured by flow cytometry from 3-to-35 d.p.t. following transient transfection of dCasMINI-modulator fusions and targeting sgRNA plasmids. Super-activation of EF1ɑ-GFP by vIRF2 screen tile (blue) is sustained for 29 days, while suppression is sustained to varying degrees by KRAB, KRAB-DNMT3L-DNMT3A, and a subset of viral and archaeal modulators (colors). Relative dCas-modulator-mCherry expression is shown (broken red line) to indicate presence of modulator plasmid expression at each time point.

**Extended Data Fig. 9.**
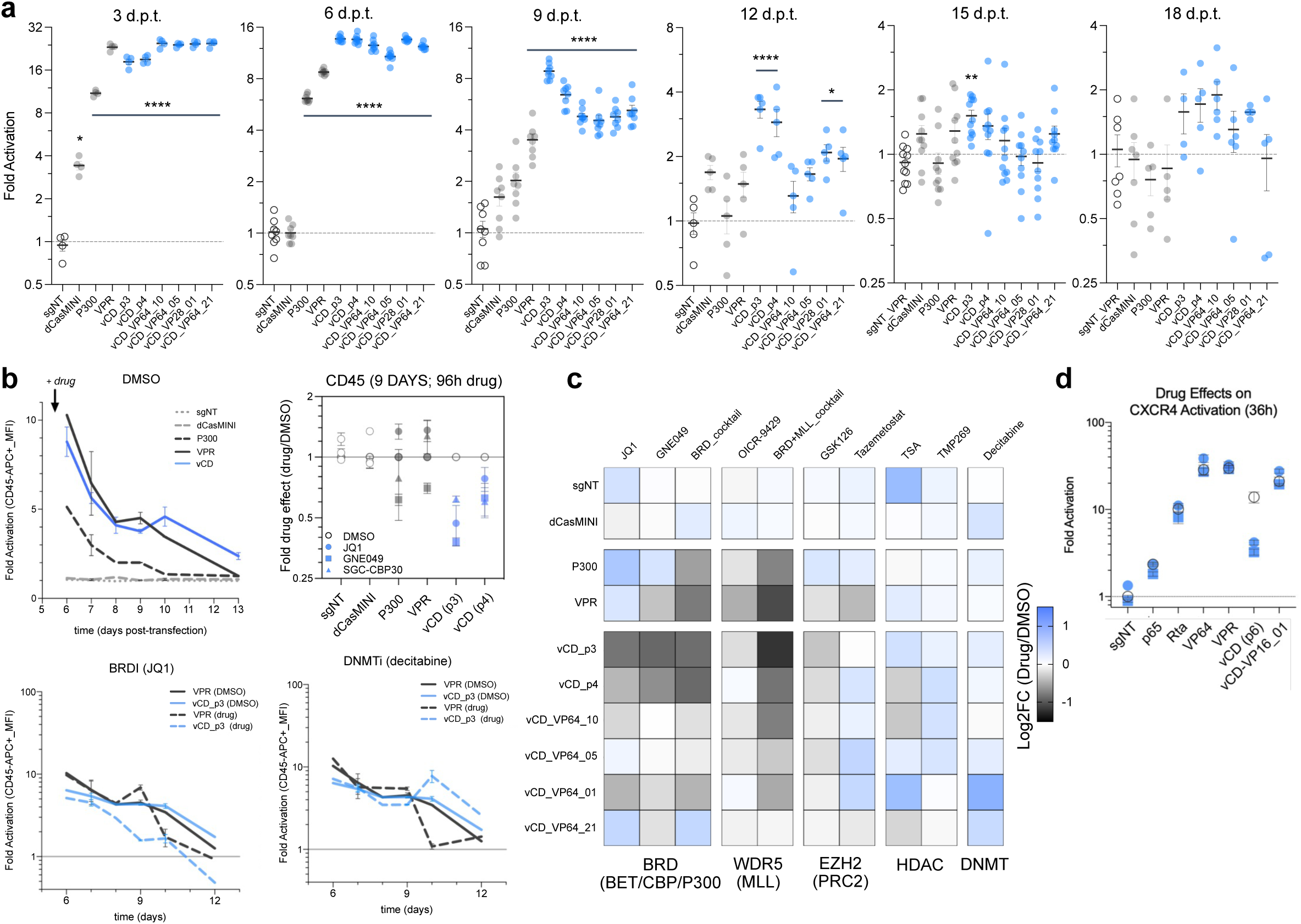
Bromodomain dependence of vCD activator durability. **a,** Independently repeated CD45 experiment (as in Fig. 4h) testing selected modulators from 3-18 d.p.t. in the presence of selective epigenetic inhibitor drugs starting from 5 d.p.t. Observed activation levels in DMSO vehicle-treated control conditions for each modulator (mean±SEM, significance from sgNT, ****P<0.0001, *P<0.05, one-way ANOVA). **b,** Observed CD45 activation levels in DMSO vehicle-treated control conditions (solid lines) for VPR (gray) and vCD_p3 (blue) (mean±SEM), or in the presence of drugs (dashed lines) BET bromodomain inhibitor JQ1 or decitabine. Normalized drug effects (top right) on CD45 activation at 9 d.p.t. (mean±SEM) of indicated vCD and benchmark modulators in the presence of selective epigenetic inhibitor drugs. **c,** Heatmap summary of normalized drug effects on CD45 activation at 9 d.p.t. of indicated vCD and benchmark modulators in the presence of selective epigenetic inhibitor drugs (**Supplementary Fig. 1**). Observed mean activation levels in DMSO vehicle-treated control (n=8) served as normalization denominators to quantify drug effects on CD45-APC fluorescence intensity. Modulator-dependent abrogation of CD45 activation in vCD-transfected cells in the presence of either BET bromodomain inhibitor alone, JQ1 or GNE049, and non-modulator-dependent abrogation of CD45 activation in vCD-, VPR-, and P300-transfected cells in the presence of equimolar cocktail of JQ1, GNE049, and histone H3K4 methyltransferase (MLL) inhibitor OICR-9429. **d,** CXCR4 activation by vCD_p6, vCD-VP16 fusion, and controls at 3-d.p.t. (mean±SEM) in the presence of BET bromodomain inhibitors JQ1 or GNE049 (blue) or DMSO-treated control (empty).

**Extended Data Fig. 10.**
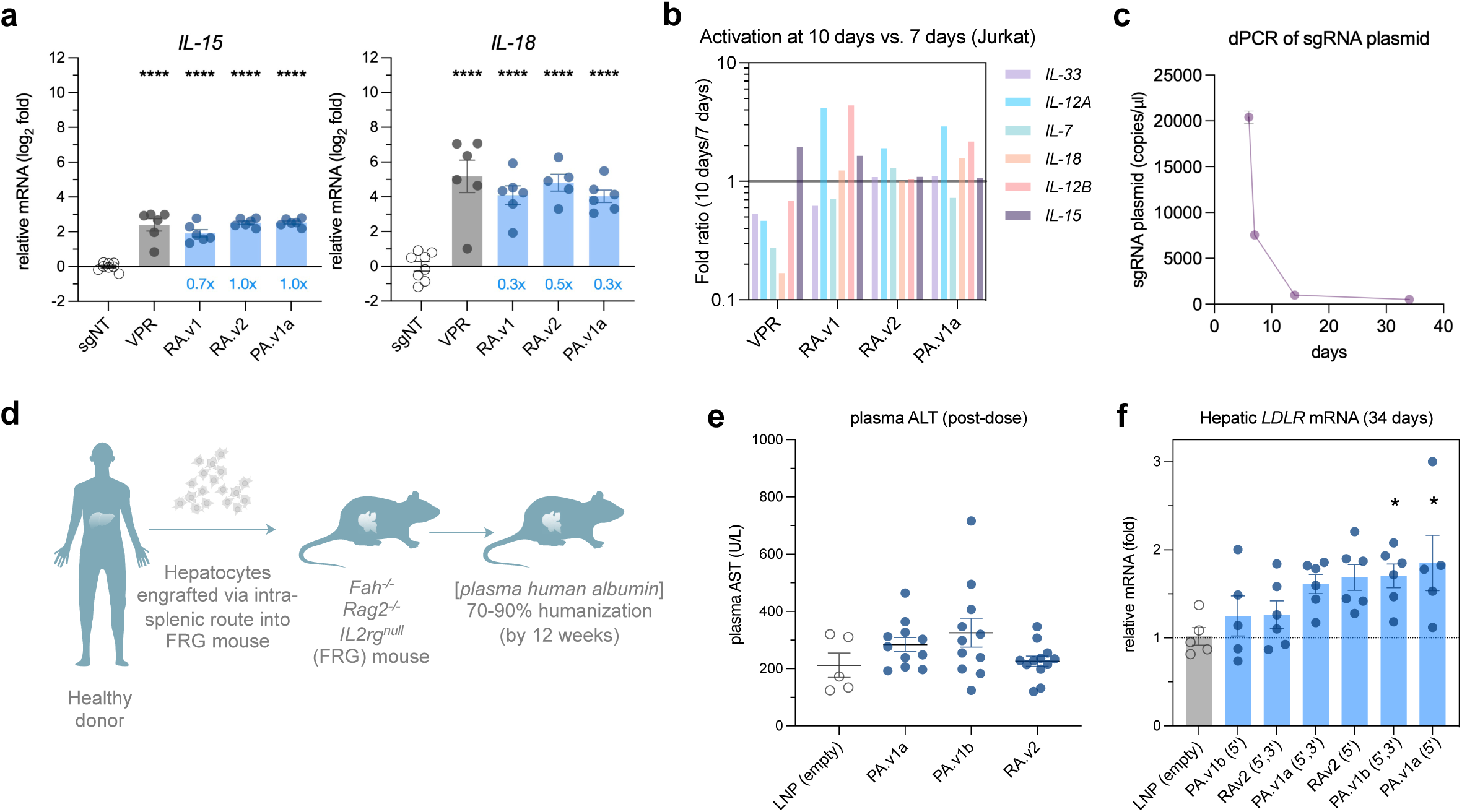
Persistent activation of therapeutically relevant genes *in vitro* and *in vivo*. **a,** *IL-15* and *IL-18* activation in dCas-modulator expressing Jurkat cells as measured by RT-qPCR, following sgRNA delivery by lentiviral transduction at 7- and 10-days post-transduction (d.p.t.). Relative mRNA fold changes, normalized to non-targeting sgRNA conditions, are plotted as mean±SEM of 3 replicates, significance from sgNT, ***P<0.0001, **P<0.001, *P<0.01, one-way ANOVA. **b,** Plotted ratios comparing activation levels at 10 days versus 7 days following lentiviral delivery (Fig. 5c**, Extended Data Fig. 10a**). **c,** digital PCR (dPCR) analysis of sgRNA copy numbers in Jurkat cells transiently nucleofected for the time series *IL-21* activation experiment (shown in Fig. 5c). **d,** Schematic depiction of the liver humanization strategy in *Fah^-/-^, Rag2^-/-^, IL2rg^null^* (FRG) mice for the vCD-mediated *LDLR* activation *in vivo* experiment. **e,** Quantification of 34-day post-dose measurements obtained from experimental mice for serum alanine aminotransferase (ALT). **f,** RT-qPCR measurement of hepatic *LDLR* mRNA sampled 34-day post-dose (as in Fig. 6h), stratified by mRNA modification type. LNPs for each dCas-activator were encapsulated to contain RNAs having either 5’ modification only, or having both 5’ and 3’ modifications, and both conditions were tested *in vivo* for each dCas-activator construct (n=6 mice per group).

**Supplementary Fig. 1.**
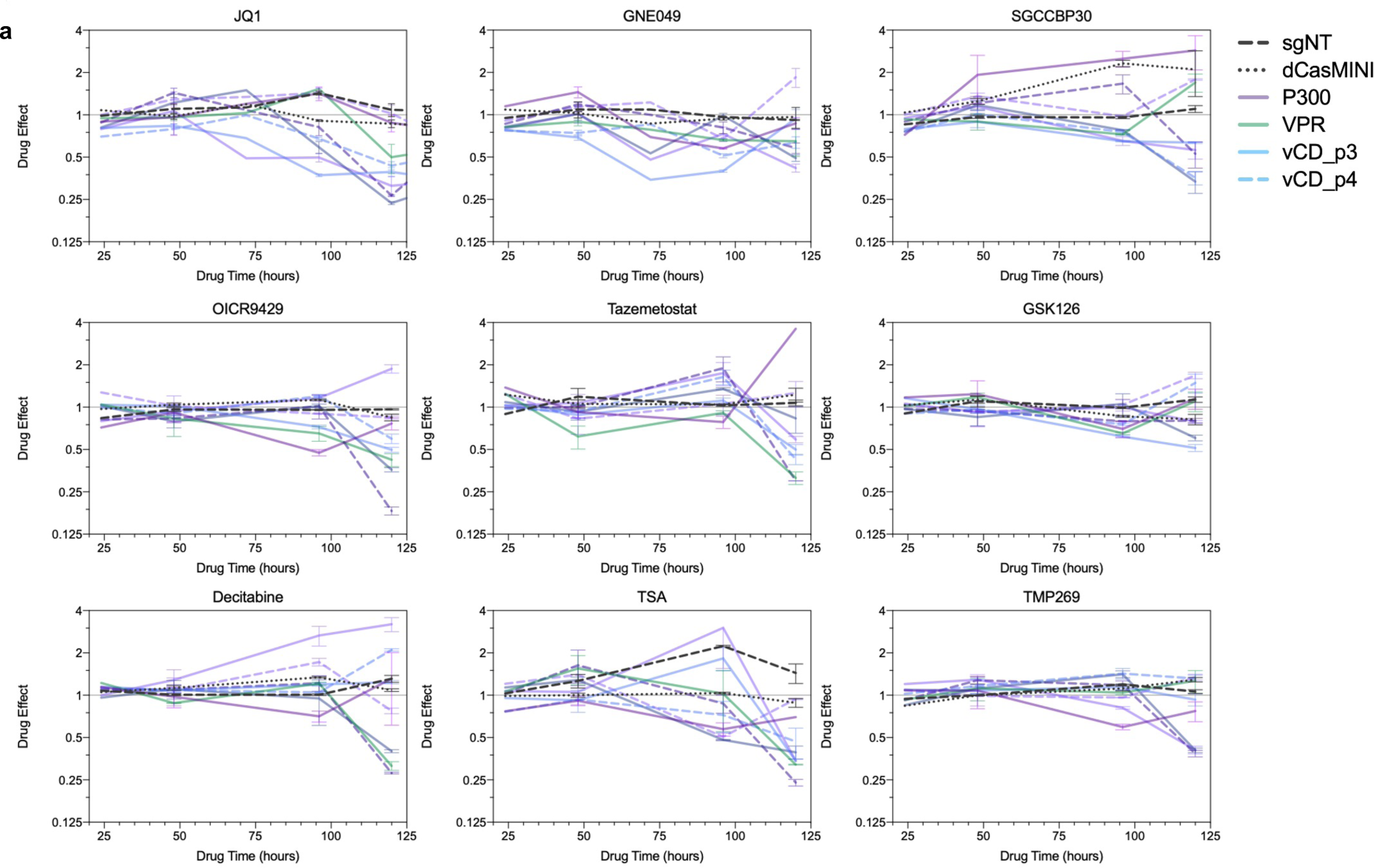
Modulator-dependent effects of epigenetic inhibitors. **a,** Normalized drug effects (mean±SEM) on CD45 activation from 6 to 10 d.p.t. of indicated vCD and benchmark modulators and negative controls, with selective epigenetic inhibitor drugs applied at 5 d.p.t. and re-dosed in fresh media each day. Observed mean activation levels in DMSO vehicle-treated control served as normalization denominators to quantify drug effects on CD45-APC fluorescence intensity (y-axis). Small molecule inhibitor drugs were chosen to selectively target BET/BRD4i/CBP/P300-associated bromodomains (JQ1, GNE049, SGC-CBP30), MLL/WDR5 and EZH2 histone methyltransferases (OICR-9229, tazemetostat), histone deacetylases (TSA, TMP269, RG2833), and DNA methyltransferase (5-aza-cytidine analog decitabine).

**Supplementary Fig. 2.**
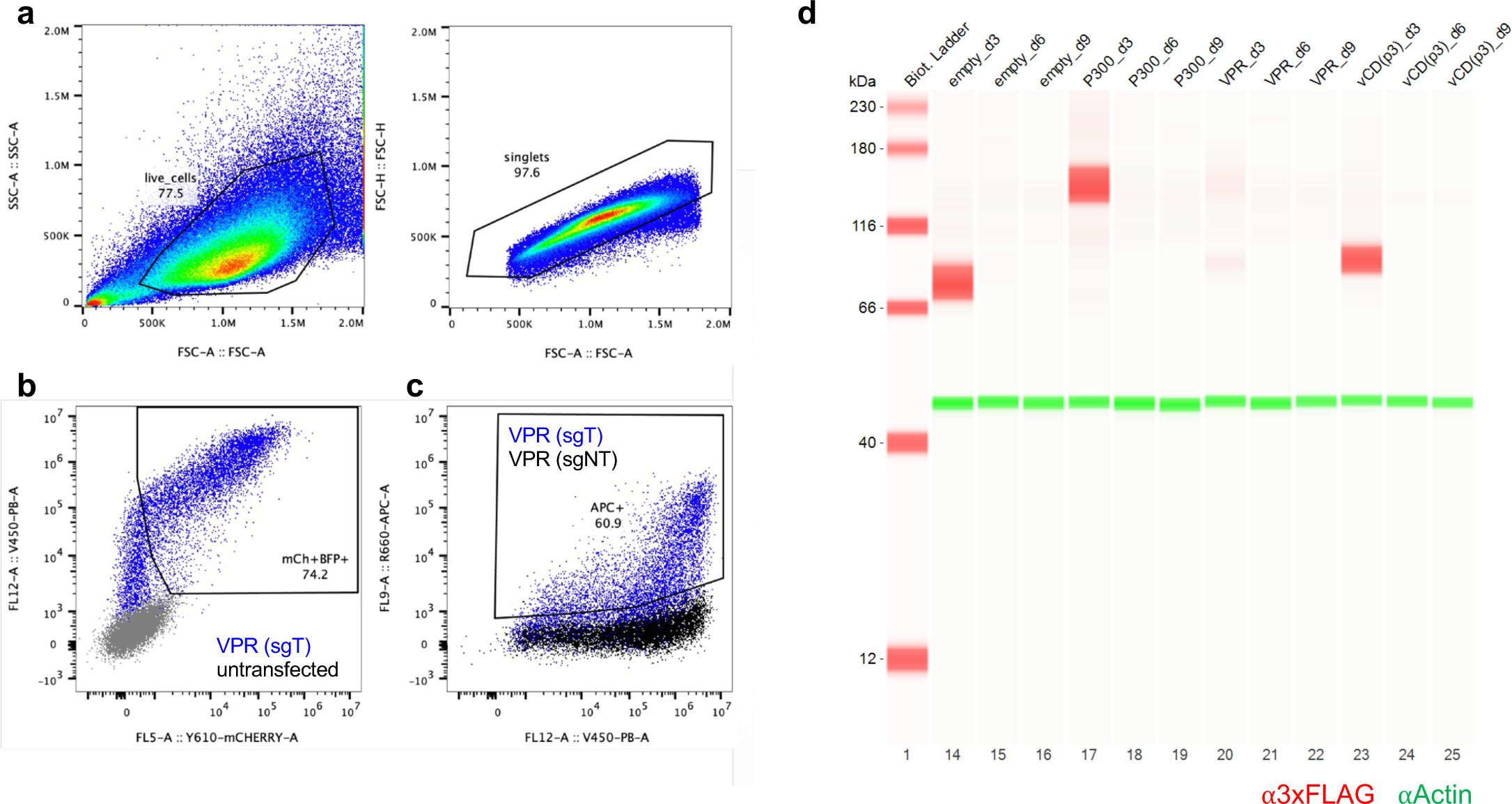
Flow cytometry gating and Western blots. **a,** Representative flow cytometry gating strategies to select live, singlet HEK293T cells. **b**, quantification of cells transfected with dCasMINI-modulator-IRES-mCherry plasmid (x-axis) and sgRNA-BFP plasmid (y-axis), compared to untransfected control (grey). **c**, representative gating of CD45-APC+ cells at 3 d.p.t. following transient transfection of dCasMINI-VPR with either targeting sgRNAs (blue) or non-targeting sgRNAs (black). **d**, Uncropped images of Western blots for detection of 3XFLAG-tagged dCasMINI-modulator-IRES-mCherry plasmid at 3-, 6-, and 9-d.p.t. for indicated modulators following transient transfection in HEK293T cells.

## Supplementary Information

Supplementary Fig. 1 | Modulator-dependent effects of epigenetic inhibitors

Supplementary Fig. 2 | Flow cytometry gating and Western blots

Supplementary Table 1: Full library screen tiles, data, and biochemical features

Supplementary Table 2: Validation sub-library screen tiles, data, and biochemical features

Supplementary Table 3: Sequences and individual recruitment data for viral and archaeal tiles in HEK293T cells

Supplementary Table 4: Sequences and individual recruitment data for engineered modulators in HEK293T cells

## Methods

### Cell lines

Experiments were carried out in K562 cells (ATCC; CCL-243), HEK293T cells (ATCC; CRL-3216), Jurkat clone E6-1 cells (ATCC; TIB-152), or HepG2 cells (ATCC; HB-8065). Cells were cultured in a humidified incubator at 37℃ and 5% CO_2_, in either RPMI 1640 (Gibco, 61870036) media (K562 and Jurkat cells) or DMEM (Gibco, 10569010) (HEK293T and HepG2 cells), supplemented with 10% FBS (Takara 632180). LentiX cells (Takara 632180) were used to produce lentivirus. The custom GFP reporter construct was generated by cloning 7 copies of a synthetic guide RNA recognition sequence (5’-TTTA GTTGTTCTAAACGCTCTGAG CGG-3’), with CasMINI and Cas9 PAM sequences, upstream of a minimal CMV promoter (miniCMV) or constitutive human EF1ɑ promoter followed by a Puromycin resistance cassette and EGFP with an intervening P2A self-cleaving peptide (Puro-P2A-EGFP) in a lentiviral transfer plasmid. These constructs were packaged into lentivirus and used to transduce K562 cells. After recovery, transduced activation reporter cells were enriched by puromycin selection (1µg mL^-1^) following nucleofection of dCas9-VPR mRNA (TriLink) and a Cas9 sgRNA targeting the ESR protospacer (IDT). Suppression reporter cells were enriched by puromycin selection directly. Single cells of each reporter cell line were isolated by serial dilution in 96-well plates. After expansion, individual clones were validated by co-transduction of dCas9-VPR or dCas9-KRAB and ESR sgRNA lentivirus and analyzed by flow cytometry to determine the dynamic range of GFP expression for each clone following activation or suppression by dCas9-VPR or -KRAB. Top performing clones were selected for further expansion and used in downstream experiments.

### Tiled library design and cloning

Human nuclear factors were determined by the searching Human Protein Atlas (.org) for "subcell_location:Nucleoplasm,Nuclear speckles,Nuclear bodies AND subcell_location:Nuclear membrane,Nucleoli,Nucleoli fibrillar center NOT subcell_location:Actin filaments,Aggresome,Cell Junctions,Centriolar satellite,Centrosome,Cleavage furrow,Cytokinetic bridge,Cytoplasmic bodies,Cytosol,Endoplasmic reticulum,Endosomes,Focal adhesion sites,Golgi apparatus,Intermediate filaments,Lipid droplets,Lysosomes,Microtubule ends,Microtubules,Midbody,Midbody ring,Mitochondria,Mitotic spindle,Peroxisomes,Plasma membrane,Rods". Sequences encoding human viral transcriptional regulators (hvTRs) were obtained from published sources^26^. Sequences encoding viruses of the families *Adenoviridae, Arenaviridae, Bornaviridae, Coronaviridae, Filoviridae, Flaviviridae, Hepadnaviridae, Herpesviridae, Orthomyxoviridae, Papillomaviridae, Paramyxoviridae, Parvoviridae, Peribunyaviridae, Phenuiviridae, Pneumoviridae, Polyomaviridae, Poxviridae, Retroviridae,* and *Rhabdoviridae,* sequences encoding viruses with known zoonotic transmission, *Flaviviridae, Lyssaviridae, Filoviridae, Paramyxoviridae, Orthomyxoviridae, Coronaviridae, Reoviridae, Togaviridae, Phenuviridae, Hantaviridae*, *Adenoviridae, and Poxviridae,* metagenomic viruses of families *Siphoviridae, podoviridae, and myoviridae,* and archaeal genome *Acidianus infernus* were manually obtained from dBatVir and NCBI. Screened peptide tiles were encoded by DNA oligos (Twist) 300 nucleotides in length, of which 255 nucleotides were target specific. The 5’ and 3’ ends of each oligo consisted of sequences complementary to the destination vector, with the 3’ 15 nucleotide overlap being composed of part of the Illumina Read 1 Primer, where [vector overlap 1-target sequence-stop-barcode-vector overlap 2 (illumina Read 1 partial)]. This design allowed for convenient cloning of the library using either NEB HiFi or In-Fusion cloning approaches, and for the convenient downstream generation of Illumina-compatible NGS libraries. For the validation sub-library: 954 predicted activators (FDR<0.25; log_2_FC>1), 1,228 predicted suppressors (FDR<0.001; log_2_FC<-1), and 22 predicted dual-activity modulators (enriched in both activation and suppression lists) were tested alongside literature-based positive control tiles, namely published activator tiles from yeast and human transcription factors^10^, plus activator and suppressor tiles from Pfam-annotated human proteins^18^. As negative controls, we included a set of 949 peptides predicted to be inactive: 314 tiles depleted from ON bins in initial mCMV and EF1ɑ screens, 563 scrambled sequence tiles with opposing activities in initial mCMV and EF1ɑ screens, and 72 tiles with early stop codons and opposing activities in initial mCMV and EF1ɑ screens.

### Pooled library screening

The custom DNA oligo library (each 300 bp, Twist) of 43,938 putative modulator elements (original screen) and ∼3,750 elements (sub-library validation screen) paired to unique 12mer DNA barcodes was cloned into dCas9 lentiviral expression plasmid at high coverage (1,000x), packaged into lentivirus, and transduced into the mCMV and EF1ɑ GFP reporter cell lines at MOI=0.3. Cells were treated with blasticidin (10µg mL^-1^) to enrich for positively transduced cells, followed by fluorescence-activated cell sorting (BD FACSAria) to separate populations of interest: mCMV-GFP-ON cells for activation and EF1ɑ-GFP-OFF cells for suppression, at 6- and 10-days post-transduction, respectively. Sorted populations of interest were further enriched by culturing for 6 additional days and subjected to 4-way gated FACS separation into discrete bins based on GFP fluorescence intensity. Genomic DNA was extracted from sorted cells in each discrete bin, and from bulk mCMV-GFP-ON and EF1ɑ-GFP-OFF cells. Barcoded modulator sequences were PCR amplified from these gDNA samples with primers containing Illumina adapter sequences and a unique i7 index for each sample. Pooled libraries were sequenced on an Illumina NextSeq 550 (Gladstone Genomics Core) to identify barcodes present in each sample.

### Pooled library screen analysis

Read count matrices for each library were generated based on alignment of sequenced modulator barcodes using a custom Python script. All subsequent data analysis was performed using R version 4.1.0. For the activation screen, technical replicates for GFP-OFF libraries were collapsed (counts per barcode were summed) resulting in two GFP-OFF replicates. For the GFP-ON conditions, we used one GFP-ON library collected at 6 days post-transduction and another that we built *in silico* by taking the weighted sum of binned GFP-ON gates P7, P8, P9, and P10 (collected after a further 6 days of enrichment). DESEQ2 (version 1.32.0) was used to identify statistically significant activator sequences that were enriched in the two GFP-ON libraries compared to the two GFP-OFF libraries (FDR<0.05, log_2_FC>0). For the suppression screen, technical replicates for GFP-ON libraries were collapsed (counts per barcode were summed) resulting in two GFP-ON replicates. For the GFP-OFF conditions, we used one GFP-OFF library collected at 10 days post-transduction and another that we built *in silico* by taking the weighted sum of GFP-OFF gates P6, P7, P8, and P9 (collected after a further 6 days of enrichment). DESEQ2 (version 1.32.0) was used to identify statistically significant suppressor sequences that were enriched in the two GFP-OFF libraries compared to the two GFP-ON libraries (FDR<0.001, log_2_FC<0). The following packages were used for downstream annotation and analysis: parSeqSim for all-by-all sequence homology to identify clusters of tiles; DECIPHER for multiple sequence alignment to identify conserved domains; DagLogo for amino acid-level enrichment of biochemical properties in tiles; MOTIF Search for enrichment of amino acid motifs and domains.

### Generalized linear regression for activator strength prediction

To identify biochemical features that were predictive of activation sequences, we trained generalized linear regression models based on the proportion of amino acids in the 85 amino acid peptide tiles (OneHot encoding) using “caret” (v6.0) and “glmnet” (v4.1) in R (v4.1.0). The top model was selected with 10-fold cross-validation and feature importance was extracted and plotted for visualization. We additionally trained generalized linear regression models based on sets of biochemical features ("biochem_tiny","biochem_small","biochem_aliphatic","biochem_aromatic","biochem_nonpolar","biochem_polar","biochem_cha rged","biochem_basic","biochem_acidic") generated using the “Peptides” package (v2.4.4).

### Statistical metrics for activation time series analysis

Normalized potency, robustness, and durability scores are defined as follows. We first calculated the average of the mean fold change per modulator, relative to baselines observed in non-targeting sgRNA (sgNT) and dCasMINI recruited without modulator fusion (dCasMINI) conditions, across replicates for the gene of interest (CXCR4, CD45, IFNG, CD81) at days 3 and 6 to define early activation potency. These averages were then plotted as mean_fold_change vs. time (days). For simplicity, we assume that the behavior of the change between time points was linear. The area under the curve (AUC) for this plot was then calculated numerically, and the subsequent errors were propagated. The AUC was then normalized to the maximum AUC for that particular gene of interest. For example, if a modulator had the highest AUC measured for CXCR4, then the normalized CXCR4 potency score for this modulator will have a value of 1. Normalized robustness score is calculated similarly, factoring modulator activity across multiple targets. Normalized durability score is calculated similarly, but the area under the curve is calculated at post-plasmid time points, i.e. all time points after day 9 post-transfection.

### Protein structure predictions

Protein structures were predicted using ESMFold^53^ within the ColabFold suite^64^. The YASARA software package^54^ was used to calculate structural features such as solvent accessible surface area and electrostatic potential. The electrostatic potential was calculated using the AMBER14 forcefield as implemented in YASARA^63^. Structural alignments were calculated using the SHEBA algorithm further checked with MUSTANG^65^. For presentation of the modulator sequence, the modulator was spaced from the Cas protein within YASARA. The modulator linker was then used to link the modulator region to the Cas protein using YASARA’s BuildLoop protocol.

### Arrayed testing of individual modulators

Oligos encoding putative human, viral, archaeal, and engineered modulator domains, and target-specific duplexed sgRNA spacer sequences, were synthesized as eBlocks (IDT) and cloned as direct fusions into dCasMINI or dCas9 vector plasmids, or sgRNA vector plasmids, respectively. For transient plasmid delivery experiments, wild-type HEK293T cells were seeded in 96-well plates at a density of 20,000 cells per well. The same day, cells were co-transfected (Mirus LT1 or Mirus X2) with uniform masses of dCasMINI-modulator- and sgRNA-expressing plasmids such that each well received a single modulator construct to be tested, but the same targeting sgRNA across all wells. Each well received 100ng of modulator plasmid and 33.3ng sgRNA plasmid and experiments were performed in triplicate at a minimum. Epigenetic inhibitor drugs were from Selleck, MedChemExpress, or MilliporeSigma.

### Flow cytometry, ELISA, RT-qPCR, Western blots

Cells were dissociated (TrypLE), washed and resuspended in flow buffer (PBS with 10% FBS and 2mM EDTA), and immunostained for surface protein detection with APC direct-conjugated primary antibodies against CD45, CXCR4, CD81, CD2, CD71 (BioLegend; 1:100 dilution) unless intended for integrated GFP reporter expression. Cells were analyzed by flow cytometry (Cytoflex LX or Attune CytPix) with analysis gates (FlowJo) to ensure measurements of live, singlet, and double-transfected cells to verify both dCas-modulator and sgRNA plasmid expression via mCherry and BFP fluorescence, respectively. Geometric mean of APC fluorescence or GFP, or percent CD45-APC+ frequency of parent for each condition were normalized against those of negative controls and reported as fold-change relative to negative controls. For IFNG ELISA assay, cell supernatants were collected to monitor IFNG protein expression by ELISA according to manufacturer protocols (BioLegend). Antibodies used: BioLegend CXCR4-APC (#306510), BioLegend CD45-APC (#304037), BioLegend CD81-APC (#349510), BioLegend CD2-APC (#309224), BioLegend CD71-APC (#334108), BioLegend B2M-APC (#395712), Miltenyi IFNG (#130-113-497), Miltenyi CD2 FITC (#170-081-028), BioLegend IFNG ELISAMAX (#430104), BioLegend CD29-APC (#303008), BioLegend CD151-APC (#350406). For cell cycle assays, cells were pulsed with 1µM 5-ethynyl-2’-deoxyuridine (EdU) for 1 hour to label S-phase cells, followed by PBS wash and replacement of fresh DMEM/FBS media. EdU detection was by Click chemistry-based AlexaFluor-488 flow cytometry assay (ThermoFisher). For RT-qPCR, cell pellets were flash-frozen and stored in -80℃ before processing for cell lysis and analysis by Cells-to-CT 1-Step TaqMan Kit with Taqman probes (ThermoFisher). For Western blots, transfected HEK293T cells were harvested at the indicated time points and resuspended in ice-cold RIPA buffer with protease inhibitor cocktail (Roche). Total protein was quantified by BCA assay (Pierce). Western blot was performed using the Jess 12-230kD separation module (ProteinSimple; SM-W001) following the manufacturer’s protocol. Lysates were normalized and diluted in 0.1x sample buffer to ∼1µg/µl, then combined with 5x Fluorescent Master Mix. Samples were denatured 5’ at 95°C and 3µl each sample was transferred to individual wells of the detection module plate. A multiplex primary antibody mix was prepared in milk-free antibody diluent using Rabbit anti-DYKDDDK antibody (R&D Systems MAB8529; 1:1200) and mouse anti-Beta actin (R&D Systems MAB8929; 1:50). Anti-rabbit-NIR and anti-mouse-IR secondary antibodies (ProteinSimple, DM-007 & DM-010) were used at 1:20 dilution in milk-free antibody diluent.

### Modulator potency assays in Jurkat cells

Lentiviruses carrying transgenes for dCas-vCD_p3, dCas-vCD_VP64, dCas-vCD_VP16, and dCas-VPR and mCherry & blasticidin resistance reporters were generated and then transduced at <0.3 MOI into Jurkat Clone E6.1 (ATCC) in arrayed fashion, resulting in stably expressing modulator cell lines. Post blasticidin selection at 20ug mL-1 (InvivoGen, ant-bl-05) stable modulator Jurkat cells were transduced with lentiviruses encoding puromycin resistance and guide protospacer sequences targeting human *IL-7, IL-15, IL-18, IL-21, IL-12a, IL-12b, and IL-33*. A non-human protospacer sequence delivered by lentivirus was used as a non-targeting control. Following puromycin selection at 2ug mL-1 (InvivoGen, ant-pr-1), the jurkat cells were harvested at several different timepoints ranging from 7 days to 14 days. At each harvest, with the reagents provided by TaqMan Gene Expression Cells-to-CT Kit (Invitrogen, AM1728), cells were lysed, then bulk RNA converted to cDNA through RT reactions, and afterwards specific cDNA transcript sequences of the above interleukin gene panel amplified by TaqMan probe based qPCR reactions (ThermoFisher, 4331182). Per qPCR reaction, multiplexing of the gene of interest and the housekeeping gene RPL19 was performed, with the gene-of-interest detected by a FAM dye and RPL19 by VIC dye. Three technical replicates were run per qPCR reaction and delta delta Ct analysis executed to calculate gene-of-interest mRNA fold change against the non-targeting controls.

### Modulator durability assays in Jurkat cells

Lentivirus carrying transgenes for dCas-vCD_p3 and dCas-VPR (mCherry & blasticidin resistance reporters included in the same constructs) were generated and then transduced at <0.3 MOI into Jurkat Clone E6.1 (ATCC) in arrayed fashion, resulting in stably expressing modulator cell lines. Post blasticidin selection at 20µg mL-1, dCas guide plasmid pool encoding protospacer sequences for *IL-21* were delivered by nucleofection into the stable modulator Jurkat cells. A non-human protospacer sequence was used as a non-targeting control. To execute nucleofection delivery of guides, 1e6 cells were loaded across two different nucleocuvettes with SE reagent and electroporated under program CL-120 on a 4D-Nucleofector X Unit (Lonza, NC0475784). Following puromycin selection at 2µg mL-1 (InvivoGen, ant-pr-1), the nucleofected cells were harvested at several different timepoints ranging from 6 days to 41 days. At each harvest, with the reagents provided by Quick-RNA Microprep Kit (Zymo, R1051), cells were lysed to release free floating RNA which was then purified to a final elution volume. Next, bulk RNA was converted to cDNA through Maxima H Minus cDNA Synthesis reactions (Thermo Scientific, M1682). Afterwards, *IL-21* cDNA transcript sequences were amplified by TaqMan probe based qPCR reactions (ThermoFisher, 4331182) using the Applied Biosystems TaqMan Fast Advanced Master Mix (Applied Biosystems, 4444963). Per qPCR reaction, multiplexing of the gene-of-interest and the housekeeping gene RPL19 was performed, with *IL-21* detected by a FAM dye and RPL19 by VIC dye. Three technical replicate RT-qPCR reactions were run per biological replicate and delta delta Ct analysis executed to calculate *IL-21* mRNA fold change against the non-targeting controls. Guide expression was measured from RNA samples derived from post-nucleofection days 6, 7, 14, and 34 through Qiacuity digital PCR (Qiagen, 250131).

### Synthetic mRNA, gRNA, and LNP generation

mRNA and gRNA were synthesized and packaged in LNP with ionizable lipid ALC0315 by Genscript. Briefly, mRNA sequences containing dCas-Activators were synthesized with a cap at the 5’ end and 100 Poly-A tail at the 3’ end along with N1-Me-nucleoside triphosphate modifications. Five unique sgRNAs targeting human LDLR were synthesized with 2’OMe and phosphorothioate modifications at the 5’ and 3’ ends, respectively, and separately with 2’OMe at the 5’ end only. Respective mRNAs and multiplexed sgRNAs were packaged in ALC0315 LNP at a 1:1 wt/wt ratio. LNP #1 (MC3), LNP #2 (SM102), LNP #3 (ALC0315) and LNP #4 (LP01) encapsulating f-luciferase mRNA also were provided by Genscript.

### Generation of humanized-liver mouse models

PXB mice and *Fah^-/-^, Rag2^-/-^, ILR2RG^-/-^*(FRG) mice were generated by Phoenix Bio and Yecuris Corporation respectively, as previously described^72^. Briefly, uPA/SCID mice (Phoenix Bio) and FRG mice (Yecuris Corporation) were injected intra-splenically with healthy human hepatocytes, which ultimately form a fully reconstituted humanized liver in 8-10 weeks. In both mouse models, humanization of the liver was determined by measuring human albumin levels in serum.

### Animal care

All animal studies were performed in accordance with the Institutional Animal Care and Use Committee Policy of Yecuris corporation (Seattle, WA) in full compliance with all relevant ethical guidelines, standard operating procedures and regulations. PXB and FRG mice were respectively housed at CRADL (South San Francisco, CA) and Yecuris corporation (Seattle, WA). General procedures for animal handling care, housing and euthanasia at both facilities were approved by the Institutional Animal Care and Utilization Committee (IACUC). PXB mice were given irradiated PROLAB ISOPRO RMH 3000 feed and drinking water supplemented with L-Ascorbic acid (Cat # 25564, Sigma-Aldrich) at 0.3 mg/ml. FRG mice were given irradiated mouse chow with a low tyrosine content (0.53% PicoLab® High Energy Mouse Diet, 5LJ5 chow) and drinking water *ad libitum*. They were also administered the prophylactic antibiotic sulfamethoxazole and trimethoprim (SMX-TMP, 80 mg/mL, 16 mg/mL) every other week in their drinking water along with NTBC for three days once every month, as well as two days prior to NTBC administration before enrollment in the study. Cages were changed every two weeks at both facilities.

### D-luc LNP bioavailability and *LDLR* activation in humanized liver mice

PXB mice were dosed with four different LNP formulations LNP#1 (MC3), LNP#2 (SM102), LNP#3 (ALC-0315) and LNP#4 (LP01) encapsulating F-Luciferase mRNA (n=1/LNP) at 0.3 mg/kg by retro-orbital injection. 4, 24, 48, 72 and 96 hours later, mice were dosed with 5 μg of D-luciferin by intra-peritoneal injection and imaged by In Vivo Imaging System (IVIS). Across all five time-points, LNP#3 gave the brightest luminescence signal and was therefore selected for delivery in the *in vivo* study. For the 34 day-long *in vivo* study, FRG mice were bled at day -7 to determine plasma total cholesterol (TC) and LDL-cholesterol (LDL-C), and weighed. At day -3, mice were randomized by plasma levels of human albumin, total cholesterol (TC), LDL-C, and body weights and divided into 4 groups (n=6 mice per group). Each group was dosed with Empty LNP (control group), LNP-vCD_VP64, LNP-vCD_p3 and LNP-vCD_p4 respectively at 2 mg/kg total RNA. Intermediate body weights were measured at day 6 and day 20. At day 34, mice were euthanized and harvested for liver tissue and blood.

### RNA extraction, cDNA synthesis, RT-qPCR and dPCR from mouse liver

25 mg of liver tissue was used for RNA extraction. Tissue was homogenized using metal PowerBead tubes and QIAshredder columns (Qiagen). Total RNA was extracted from the homogenate using the PureLink RNA Mini kit (Thermo Fisher Scientific). Complementary DNA was synthesized from 500 μg of total RNA using the Maxima™ H Minus cDNA Synthesis Master Mix (Thermo Fisher Scientific). Quantitative PCR was performed on a QuantStudio 7 Flex Real-Time PCR system (Applied Biosystems). Relative gene quantification was performed using TaqMan primer probe set for LDLR (Hs01092524_m1) normalized against RPL19 (Hs02338565_gH) (Thermo Fisher). Each PCR reaction mix consisted of 0.5 μL primer probe, 5 μL of TaqMan Fast Advanced Master Mix (Thermo Fisher Scientific) and 1 μL of cDNA. Fold-changes were evaluated using the ΔΔ-Ct method. Digital PCR for dCas was conducted on a QIAcuity Digital PCR system (Qiagen). RNA was prepared according to manufacturer’s protocol using the QIAcuity OneStep Advanced Probe kit. Primer-probe sequences for dCas were as follows: forward primer (CTC ACC GAG AAA AGC GAA CG), reverse primer: (CTG GAC CGT ACC CAC TTT GT), probe (/5SUN/CTG GGC TTG /ZEN/TGA AAT TGC CGA CTT /3IABkFQ/). Samples were prepared in duplicates in a standard 96-well PCR plate and transferred to a QIAcuity 8.5k 96-well Nanoplate (Qiagen) for partitioning using the Qiagen Standard Priming Profile. Digital PCR cycling conditions were as follows: reverse transcription for 40 minutes at 50°C, RT enzyme activation for 2 minutes at 95 °C, 40 cycles of denaturation for 5 seconds at 95°C and annealing/extension for 30 seconds at 60 °C. Qiagen’s QIAcuity Software Suite was used to adjust sample thresholds.

## Data Availability

The illumina sequencing datasets generated in this study will be made available in the NCBI Sequencing Read Archive.

## Code Availability

All custom codes used for data analysis are available upon request.

## Acknowledgements

We thank Andrew Norton, Wenjun Wu, Christopher Still II, T. Danny Ko, Ian Lam, Aayushma Gautam, Mohamed Ghazal, Tabitha Tcheau, and Xiaoshu Xu for helpful conversations and assistance, and Mylinh Bernardi of the Gladstone Genomics core for assistance with illumina sequencing. L.S.Q. is supported by Sarafan ChEM-H, Stanford University and is a Chan-Zuckerberg BioHub-San Francisco Investigator.

## Author Contributions

D.O.H., L.S.Q., and T.P.D. designed the study. G.A.C., R.W.Y., D.O.H., and T.P.D. designed screening libraries. G.A.C., R.W.Y., and V.C. designed engineered activators. T.B.G. designed reporter cells and library cloning strategies. G.A.C., J.K., K.J., N.J.S, T.B.G., and V.C. performed experiments. G.A.C., R.W.Y., M.Z.J., K.J., S.V.J., A.C.D., and X.Y. analyzed data. G.A.C. and R.W.Y. wrote the manuscript with significant contributions from all authors. D.O.H. supervised the study.

